# Evidence against preserved syntactic comprehension in healthy aging

**DOI:** 10.1101/299883

**Authors:** Charlotte Poulisse, Linda Wheeldon, Katrien Segaert

**Affiliations:** School of Psychology, University of Birmingham, Birmingham, United Kingdom; Department of Foreign Languages and Translation, University of Agder, Kristiansand, Norway; Centre for Human Brain Health, University of Birmingham, Birmingham, United Kingdom

**Keywords:** healthy ageing, syntactic comprehension, processing speed, working memory

## Abstract

We investigated age-related differences in syntactic comprehension in young and older adults. Most previous research found no evidence of age-related decline in syntactic processing. We investigated elementary syntactic comprehension of minimal sentences (e.g. I cook), minimizing the influence of working memory. We also investigated the contribution of semantic processing by comparing sentences containing real verbs (e.g. I cook) versus pseudoverbs (e.g. I spuff). We measured the speed and accuracy of detecting syntactic agreement errors (e.g. I cooks, I spuffs). We found that older adults were slower and less accurate than younger adults in detecting syntactic agreement errors for both real and pseudoverb sentences, suggesting there is age-related decline in syntactic comprehension. The age-related decline in accuracy was smaller for the pseudoverb sentences, and the decline in speed was larger for the pseudoverb sentences, compared to real verb sentences. We suggest that syntactic comprehension decline is stronger in the absence of semantic information, which causes older adults to produce slower responses in order to make more accurate decisions. In line with these findings, performance for older adults was positively related to a measure of processing speed capacity. Taken together, we found evidence that elementary syntactic processing abilities decline in healthy ageing.

## Introduction

Syntactic processing is often discussed in the literature as a key example of a cognitive function that is relatively resilient to age-related decline (Campbell et al., 2016; Samu et al., 2017; Shafto & Tyler, 2014). Studies investigating the effect of age on syntactic comprehension typically use sentences with a complex syntactic structure, such as garden path sentences with a temporary syntactic ambiguity (Samu et al., 2017), or relative clause manipulations that require disambiguation of referential choices (Payne et al., 2014). The interpretation of such complex syntactic structures may not exclusively rely on syntax, but instead, may also require additional comprehension mechanisms including semantic and pragmatic processing. Consequently, such measures of complex sentence processing may not be ideal for measuring syntactic comprehension as an isolated process. Furthermore, complex syntactic structures might impose a larger burden on working memory, as long distance linguistic dependencies must be retained in working memory in order for successful syntactic and thematic integration to take place (Tan, Martin, & Van Dyke, 2017). However, for alternative views on the role of working memory in language processing, see (MacDonald & Christiansen, 2002). Given that age is associated with declines in working memory (Waters & Caplan, 2007), the use of such computationally expensive sentences is problematic. In the present work, we aim to address these issues by reducing the complexity of our stimuli to simple two word sentences, in order to investigate the comprehension of elementary syntactic structures. Consequently, contextual cues and working memory load are kept to a minimum. Moreover, we compare these elementary syntactic operations in real word versus pseudoword sentences, in order to investigate the contribution of meaning to syntactic comprehension. Lastly, we investigate whether individual differences in working memory, processing speed and physical health impact on decline in syntactic comprehension in healthy ageing.

### Syntactic comprehension

Syntax plays a fundamental role in understanding spoken language. Syntactic information, in addition to other types of information, enables the listener to extract meaning from the incoming speech input. Syntactic processes are used in sentence comprehension in a number of ways, including structure building (e.g. combining words into larger units based on grammar rules and word category information) and checking agreement (e.g. in English, the verb needs to agree in number and person with the subject) (Kaan & Swaab, 2002). Furthermore, syntax plays an important role in mapping thematic roles (e.g. mapping the agent (‘doer’) and patient (‘doe-ee’) onto certain positions in the sentence). The order of noun phrases to thematic role mapping strongly influences the complexity of the sentence structure and the number of syntactic operations needed to determine the meaning of a sentence. In sum, the level of syntactic processing required to understand spoken language can range from rather simple to very complex.

A considerable amount of research has focused on whether there is age-related decline in sentence comprehension. The emphasis in this line of research tends to be on complex sentence structures. Using a paradigm that capitalizes on syntactic ambiguity, Tyler and colleagues investigated syntactic processing during sentence comprehension in younger and older adults, in sentences varying in the level of syntactic processing required (Campbell et al., 2016; Davis, Zhuang, Wright, & Tyler, 2014; Meunier, Stamatakis, & Tyler, 2014; Samu et al., 2017; Shafto & Tyler, 2014). Specifically, unambiguous sentences have only one possible syntactic interpretation (e.g. *…“sneering boys”*), whereas ambiguous sentences have two possible interpretations: an interpretation that, given its higher frequency in the language, is dominant or more expected (e.g *“cooking apples are”*), or an interpretation that is subordinate, or less expected (e.g. *…“cooking apples is”*). Participants are asked to indicate whether the disambiguating word (*are* or *is* in the examples) is an acceptable or an unacceptable continuation of the sentence. For individuals without any language disorders, a conventional pattern of responding is to reject more (and respond more slowly to) subordinate sentences compared to dominant and unambiguous sentences, with little difference between the latter two sentences (Campbell et al., 2016). Tyler and colleagues repeatedly found no age-related differences in acceptability ratings (Davis et al., 2014; Meunier et al., 2014), or response times, in which the mean response time difference between the sentences requiring the most and the least syntactic processing (subordinate and unambiguous sentences) was used (Campbell et al., 2016; Samu et al., 2017). Another line of research has measured online syntactic processing with a word-monitoring task to investigate younger and older adults’ ability to develop syntactically and semantically coherent representations (Tyler et al., 2010). Participants listened to sentences and were instructed to press a response key whenever they heard a pre-specified target word. Word position of the target word varied from early to late across the sentences. The sentences differentially loaded on syntactic and semantic processing: normal prose sentences had a normal syntactic, semantic and pragmatic structure; anomalous prose sentences had a correct grammatical structure but lacked sentential meaning, and randomly ordered word strings lacked grammatical and sentential meaning. Response times increased at later word positions in both normal and anomalous prose. Comparing a group of young and older adults, this pattern of word position effects showed no age-related performance differences. Taken together, these results suggest that syntactic comprehension is preserved in the late years of adult life. However, all these studies have placed complex syntactic structures at the forefront. Since the manipulations in these studies potentially do not exclusively investigate the contribution of syntactic processes, it is unclear to what extent the performance for processing these sentences also reflects additional (linguistic and pragmatic) comprehension mechanisms.

Moreover, even though in a large number of studies it is concluded that syntactic comprehension performance is preserved in healthy ageing, there are also several studies that have found age-related syntactic comprehension decline. Specifically, older adults tend to be less accurate and slower in answering comprehension questions for syntactically ambiguous sentences (Waters & Caplan, 2001 and Kemtes & Kemper, 1997). Obler, Fein, Nicholas, & Albert (1991) investigated age-related differences in the effect of syntactic complexity and semantic plausibility on sentence comprehension. Participants listened to sentences that were divided into six different syntactic types (active, passive, single negative, double negative, double embedded or comparative). Accuracy showed a general age-related decline and older adults were disproportionally less accurate at the harder sentence types. In a sentence picture matching paradigm with sentences of increasing syntactic complexity, Antonenko et al. (2013) found superior syntactic performance in younger compared to older adults. The paradigm consisted of sentences with three different levels of syntactic complexity. The easiest level did not have hierarchical embeddings (e.g. *“The tiger is crying, pulling the frog, and he is gray.”*), while the other two levels included one or two subordinate clauses (e.g. *“The tiger that is crying and pulling the frog is gray/”* and *“The tiger that is pulling the frog that is crying is gray.”*). A correct picture matching decision required full understanding of the sentence structure. Older adults were less accurate and slower than younger adults in the task, but the effect of syntactic complexity was not different between age groups. The behavioural results were related to brain function and structure. Syntactic abilities of young adults were associated with functional coupling in a dedicated, mainly left hemispheric syntax network. In contrast, the syntax network of the older adults included additional (frontal and parietal) regions supporting working memory as well as semantic processing. Indeed, numerous functional imaging studies have shown that older adults recruit different, or additional brain regions compared to younger adults to perform certain tasks, with some research suggesting these additional activity patterns are compensatory in nature (Cabeza, Anderson, Locantore, & McIntosh, 2002; Grossman et al., 2002). Crucially, the finding by Antonenko et. al (2003) that syntactic ability in older adults was related to the recruitment of regions supporting working memory as well as semantic processes emphasizes the relevance of a behavioural measure that isolates the syntactic component in sentence comprehension.

### The influence of semantic processing on syntactic comprehension

Syntactic comprehension is strongly influenced by semantic information. However, there exists debate with respect to the time course within which the integration of syntactic and semantic information takes place. Serial syntax-first models assume the language processing system initially constructs a simple syntactic structure independent of lexical-semantic information and semantic aspects are integrated at a later stage (Frazier & Fodor, 1978; Kimball, 1973). In contrast, interactive-constraint models assume syntactic and semantic processes interact at any time (Marslen-Wilson & Tyler, 1980; Taraban & Mcclelland, 1988). A third approach, the neurocognitive model of auditory sentence processing (Friederici, 2002) argues that autonomous and interactive processes coexist, but describe different processing phases during language comprehension.

Some research suggests that the interplay between syntax and semantics changes with age. Specifically, older adults rely on morpho-syntactic information to a lesser degree than young adults when other cues for sentence interpretation are available (Bates, Friederici, & Wulfeck, 1987). For example, Obler et al. (1991; mentioned above) did not only investigate age-related decline in processing syntactic complexity, they also investigated whether semantic information can aid in processing syntactically complex sentences. Sentences were either semantically plausible or implausible. Older adults were disproportionately less accurate in acceptability judgements for more syntactically complex sentence types but also for implausible sentences. The authors therefore suggested that older adults come to rely more on processing strategies that stress the plausibility of the semantics of the sentences in terms of their world knowledge rather than on a strict decoding of the syntactic structure. These results are in line with more general findings suggesting that older adults increasingly rely on semantics and world knowledge in auditory sentence processing and reading comprehension (Wingfield et al., 1994; Wingfield, 1996 and Soederberg Miller et al., 2004) as well as in other domains, such as memory (e.g. Castel, 2005; Rowe, Valderrama, Hasher, & Lenartowicz, 2006). In sum, previous findings suggest that non-syntactic components such as semantics and pragmatics facilitate syntactic comprehension and that contextual information in sentence comprehension becomes more important with age.

### The moderating effect of individual differences

Although there exists a general picture of cognitive decline in healthy aging, there is also a large amount of individual variability. In fact, the heterogeneity in performance tends to increase with age (Stones, Kozma & Hanna, 1990). As comprehensibly described in a review by Peelle (in press), an individual’s performance on a language task is not only determined by the task requirements, but also by the processing resources available to that individual. The level of resources available varies widely in older adults, with processing efficiency being determined by the person’s working memory, attention and processing speed abilities, but also by neuroanatomical features (Peelle, in press). Neuroanatomical features in turn are related not only to the person’s chronological age, but also to other factors such as the person’s aerobic fitness level (Hillman, Erickson, & Kramer, 2008; Lazarus, Lord, & Harridge, 2018). Understanding what accounts for inter-individual variability in age-related decline in cognitive tasks is therefore an important issue in aging research.

It is well known that aging is associated with decline in working memory and processing speed (Waters & Caplan, 2007); both are known also to contribute to language comprehension (Just & Carpenter, 1992; Salthouse, 1996). A study by Wingfield, Peelle & Grossman (2003) on the effects of speech rate and syntactic complexity in young and older adults established the moderating influence of processing speed on age differences in sentence comprehension. In this experiment, a group of younger and older adults heard short sentences that differed in syntactic complexity by using subject relative clauses (*“Men that assist women are helpful”*) and object relative centre embedded clauses (*“Women that men assist are helpful”*). Furthermore, speech rate was time compressed to 80%, 65%, 50% or 35% of the original speaking time, varying the processing challenge. Participants were asked to indicate whether the action was performed by either a male or female character. Accuracy was lower for the more complex object-relative clause sentences than for the easier subject-relative sentences for both age groups, with older adults showing disproportionally poorer comprehension accuracy only at accelerated speech rates. While older adults were slower than younger adults at all speech rates, older adults had disproportionately longer response times for accelerated speech rates and more complex syntactic structures. In a similar vein, a number of studies have demonstrated that the influence of working memory on sentence processing is larger among older compared to younger adults. Payne et al. (2014) found that age differences in relative clause comprehension were largely modulated by individual differences in working memory and that this influence was exaggerated among older adults. Specifically, during comprehension of sentences introducing a temporary syntactic attachment ambiguity (e.g. “*The son of the princess who scratched himself/herself in public was humiliated*”), poorer working memory in older adults was associated with increased processing time in sentences in which the reflexive pronoun referred to the object of the modifying prepositional phrase (*herself the princess*). Payne et al. (2014) suggest that with increasing age, attentional control resources in working memory are recruited at progressively lower levels of difficulty in order to maintain comprehension. These findings illustrate the importance of investigating how individual differences in working memory and processing speed contribute to age-related differences in syntactic comprehension.

Another factor that has gained increasing attention is a person’s physical health. Taking into account variability in health characteristics could explain a considerable proportion of variance that would otherwise be ascribed to age (Raz, 2009). In this context, Lara et al. (2015) have proposed a set of biomarkers of healthy aging, in which healthy aging was operationalised as preserved physical, cognitive, physiological, endocrine, immune and metabolic functions. Lifestyle variables such as regular physical activity and aerobic fitness have gained much attention in research focused on differential cognitive aging (Colcombe et al., 2004) and aerobic fitness levels have been shown to be associated with word production in healthy older adults (Segaert et al., 2018). In the current study, we will measure grip strength, because it is an established marker of a person’s physical health (Lara et al., 2015) and it has previously been related to cognitive decline (Auyeung, Lee, Kwok & Woo, 2011). We will also administer a physical activity questionnaire. Addressing the moderating influence of working memory, processing speed and physical health can leverage the predictive power of research on age differences in syntactic comprehension.

### Current study

In the current study we investigate whether there is age-related decline in syntactic comprehension. Specifically, our aims are threefold. Firstly, we aim to test whether the comprehension of elementary syntactic structure is preserved in older age. Secondly, we aim to test whether lexical-semantic content aids syntactic comprehension and whether this changes with age. Thirdly, we aim to investigate whether individual differences in working memory, processing speed and physical health modulate syntactic comprehension and moreover, whether the impact of these increases with age.

We investigate syntactic comprehension in an auditory syntactic judgement task, in a group of younger and older participants. The complexity of our stimuli is reduced to simple two word phrases consisting of a pronoun and a verb (e.g. *“l walk”*). Consequently, working memory load for processing these phrases is minimal. A similar task was used in Segaert, Mazaheri and Hagoort (2018). In the present study, lexical-semantic content is varied by using existing verbs versus pseudoverbs. A pseudoword follows the orthographic and phonological rules of a language, but has no meaning in the mental lexicon of that language. The pseudoverbs were used to create phrases of minimal semantic content (e.g. *“she ploffs”*), whereas the existing verbs were used to create semantically meaningful phrases (e.g. *“she cooks”*). The pseudoverbs and existing verbs formed two separate experimental blocks, identical in all aspects but the use of the pseudoverbs versus the real verbs. We will refer to these blocks as the “Pseudoverb” and “Real verb” block respectively. The task was to listen to the phrases and indicate whether it was morpho-syntactically correct (yes/no). In addition to accuracy, response time (RT) was measured from the start of the response screen to the button press.

To investigate the impact of individual differences on syntactic comprehension, we measured important biomarkers of healthy aging (Lara et al., 2005): physical health was assessed using strength grip and a physical activity questionnaire; cognitive functioning was assessed through a working memory, processing speed and verbal IQ measure.

We predict the following. First, in line with most previous findings of preserved syntactic comprehension in aging, we predict that performance on the Real verb phrases is equivalent for young and older adults. Second, we expect reduced performance on the Pseudoverb phrases for older adults, compared to young adults, in line with previous findings suggesting that older adults come to rely more on strategies involving semantic processing. We also expect a stronger influence of working memory and processing speed for older compared to young adults. Lastly, if a relationship exists between physical health and syntactic comprehension in older adults, we expect to find that age-related decline in syntactic comprehension is modulated by physical health, with higher levels of physical health associated with better performance in older adults.

## Methods

### Participants

50 young university undergraduates (45 women, mean age: 19, SD: 0.92, 5 men, mean age: 20 y, SD: 0.89) and 50 older adults (28 women, mean age: 71, SD: 5.79, 22 men, mean age: 72, SD: 5.68) participated in the study. Participants were recruited via the database of the School of Psychology of the University of Birmingham. All participants were native British English speakers with normal or corrected to normal hearing. Exclusion criteria included bilingualism, neurological disorders, speech or language disorders and dyslexia. To assess general cognitive function, the Montreal Cognitive Assessment test (MoCa; version 7.1) was administered to the elderly participants, resulting in 5 participants being excluded, as their scores were equal to or below the cut-off value of 26. Consequently, 45 older participants (23 women, mean age: 71, SD: 5.66 and 22 men, mean age: 73, SD: 5.61) were included in the analyses. The older participants’ education level ranged from Primary School (1 participant); O-levels/GCS2 (11); A levels/Vocational Course (6); Bachelors/Undergraduate level (21) and Master’s degree or higher (10). All participants gave informed consent. Students were given university credits as compensation; older adults received monetary compensation. The research was conducted at the University of Birmingham and had full ethical approval.

### Materials

A set of 20 English pseudoverbs created by Ullman et al. (1997) served as stimulus materials for the Pseudoverb block: brop, crog, cug, dotch, grush, plag, plam, pob, prap, prass, satch, scash, scur, slub, spuff, stoff, trab, traff, tunch, vask. These pseudoverbs were all monosyllabic with an average word length of four letters and an average phoneme length of 3.7. All pseudoverbs could be inflected according to regular grammar rules for verbs in English. They could be combined with six pronouns (I, you, he, she, we, they) or with 6 adverbs (daily, quickly, safely, early, promptly, rarely). This would yield minimal phrases, such as “I dotch”, “he dotches”, “they dotched”, or “dotch quickly”. In addition, a set of twenty common English verbs were selected to serve as stimulus material for the Real verb block: chop, cook, cram, bake, drop, flap, skip, brew, rob, rush, scour, move, jog, slam, stir, tug, walk, pull, stack, reap. These were regular monosyllabic verbs, matched in length to the pseudoverbs with an average phoneme length of 3.5. Like the pseudoverbs, these real verbs could be combined with a pronoun, or an adverb to form minimal phrases, such as “I chop”, “she chops”, “they chop”, or “chop quickly”. The same adverbs were used with both the pseudoverbs and real verbs. The adverbs were all disyllabic and care was taken to ensure that combining them with any of the real verbs would form a semantically meaningful combination.

Digital recordings of all stimuli were made using a male native speaker of English. All verbs were recorded in first, second and third singular and plural present tense. Each stimulus was pronounced three times, after which the clearest recording was selected. In order to equalize the volume of the individual recordings, all audio files in wav format were normalized to 1db using the software program Adobe Audition.

### Design

The order of the Real verb and Pseudoverb blocks was counterbalanced across participants. Both blocks consist of the same four conditions (see Table 1). In the *Correct syntax condition* a (pseudo)verb was paired with a pronoun, resulting in a morpho-syntactically correct combination (e.g. “she cugs”, “she walks”). In the *Incorrect syntax condition*, integration could be attempted, but the inflection of the verb/pseudoverb did not match the pronoun (e.g. “she cug”, “she walk”). In addition, two filler conditions were included. For the *No syntax filler condition*, the verb/pseudoverb was paired with another verb/pseudo (e.g. “dotch cugs”, “bake walks”). This combination of stimuli should not trigger integration processes at a morpho-syntactic level. The No *syntax* filler condition was included in the current experiment in order to verify that participants indeed read these phrases as a pairing of two verbs/pseudoverbs and did not attempt to integrate them. The purpose of this condition (merely a filler condition in the present experiment) was to include it as a condition of interest (a baseline condition) in a follow-up EEG experiment. Finally, the *Adverb filler condition* consisted of a verb/pseudoverb paired with an adverb (e.g. “cugs quickly”, “walks quickly”). The purpose of the Adverb fillers was to avoid any predictability in the engagement of integration processes for pairs beginning with a verb/pseudoverb. Specifically, a word pair starting with a verb/pseudoverb had an equal chance of forming a syntactically correct or incorrect word pair. To briefly preview the results, participants were highly accurate on the filler trials (above 90% across experimental blocks in both age groups), suggesting participants understood the task. An overview of the stimulus sets for both blocks and examples of all conditions is provided in Table 1.

**Table 1.** Example stimuli in each condition for the Pseudoverb and Real Verb block with trial number per condition.

| Condition [number of trials] | Example Pseudoverb block | Example Real Verb block | Correct sentence? |
| --- | --- | --- | --- |
| Correct syntax Condition [36] | I ploff<br>she ploffs<br>we ploffed | I pull<br>she pulls<br>we pulled | Yes |
| Incorrect syntax Condition [36] | I ploffs<br>he ploff | I pulls<br>he pull | No |
| No syntax filler [36] | ploffs dotch<br>ploff dotches<br>ploff dotched | walks pull<br>walk pulls<br>walk pulled | No |
| Adverb filler [36] | ploff quickly<br>ploffs quickly<br>ploffed quickly | walk quickly<br>walks quickly<br>walked quickly | Yes |

### Task

Participants were tasked with detecting grammatical mistakes. The timing of each component in one trial is illustrated in Figure 1. Each trial started with a fixation cross (1000 ms) and a blank screen (1000 ms). Following this, the minimal phrase was presented word by word with a Stimulus Onset Asynchrony of 1200 ms. The Inter Stimulus Interval (ISI) between the first and the second word varied as a function of the duration of the first word and ranged between 300 and 900 ms. A response screen showing the text “*Was this a grammatically correct sentence*?” appeared 805 ms after the onset of the second word and remained on the screen until a button press. The ISI between the second word and the response screen varied between 100 and 505 ms as a function of the duration of the second word. Participants were instructed to indicate whether the word pair they just heard was grammatically correct by clicking the left and right mouse button to respond with ‘yes’ or ‘no’ respectively. The response screen was followed by a blank screen for 6 ms. The correct response for each condition is listed in Table 1. The experiment was run using the E-Prime 2.0 software (Psychology Software Tools, Pittsburgh, PA).

**Figure 1.**
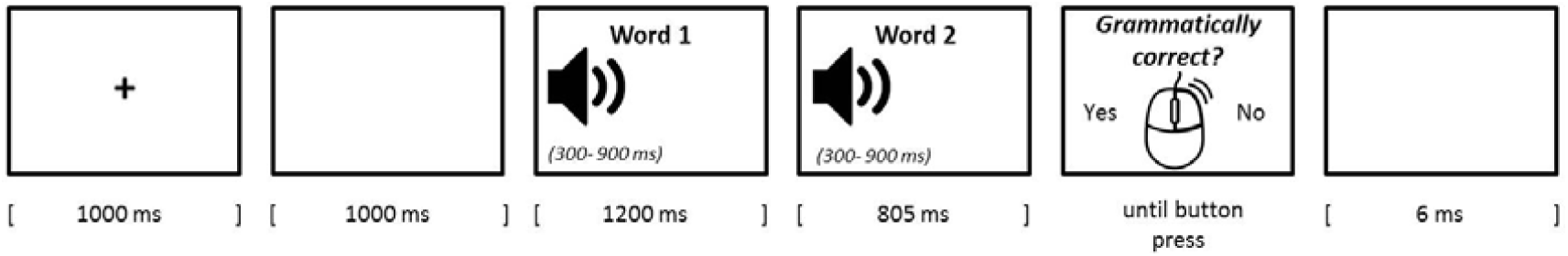
Timing of the components of one trial.

### Experimental lists

As can be seen in Table 1, the Correct syntax condition can be formed with three possible pronoun -verb/pseudoverb combinations. That is, the verb/pseudoverb *stem* combined with either ‘I’, ‘you’, ‘we’ or ‘they’; the verb/pseudoverb stem plus -*s* combined with ‘he’ or ‘she’, or the verb/pseudoverb stem plus *-ed* combined with each of the six pronouns. Each form occurred 12 times and the possible pronouns within each form occurred an equal number of times. This means that each possible pronoun occurred 3 times in the *stem form*, 6 times in the *-s form* and 2 times in the *-ed form.* The verbs/pseudoverbs were randomly assigned to the pronouns, with the constraint that each verb would occur only once in each form. The Incorrect syntax word pairs were formed according to the same criteria. However, as no incorrect combination can be composed with the *-ed form*, only two forms were possible. To ensure an equal number of trials across conditions, both the *stem form* and -s *form* consisted of 18 trials in this condition, again ensuring that the possible pronouns occurred an equal number of times. The No syntax filler condition consisted of three possible forms, such that the second verb could either be *stem-form, -s form*, or *-ed form*, with 12 trials per form. To avoid repetition effects, the first word of the pair in this condition could neither be the same verb nor have the same ending as the second word of the pair. Lastly, the Adverb filler condition also consisted of three possible forms, with the first word being either in *stem-form, -s form*, or *-ed form*, followed by randomly assigned adverbs as the second word. There were 36 trials per condition, resulting in 144 trials in total for both blocks.

A unique randomized stimulus list was created for each participant and divided into three separate sections, separated with self-paced breaks. The order of the Pseudoverb and Real Verb block was counterbalanced between participants. Each block was preceded by a unique list of 33 practice trials.

### Inter-individual variability markers

A number of individual differences measures were collected to assess the physical health and cognitive functioning of our participants.

### Markers of cognitive function

The Backward Digit Span task (Waters & Caplan, 2003) was administered to measure *working memory capacity.* Using the E-Prime 2.0 software (Psychology Software Tools, Pittsburgh, PA), participants were instructed to attend to a series of visually presented digits of increasing length. After the presentation of the last digit, participants were instructed to enter the digits in the reverse order by using the numbers on the keyboard. The task began at a length of two digits and went up to seven digits. There were 5 trials at each digit length. No practice trials were included. Span size was defined as the longest digit length at which a participant correctly recalled three out of five trials. If a participant recalled two out of five trials correctly at the longest digit length, half a point was added to the total score. The raw span size scores were used in the analyses.

Using the WISC-IV Coding subtask (WAIS-IV; Wechsler, 2008), *processing speed* was assessed. In this task, the participant is asked to copy symbols that are paired with numbers within 120 seconds. A point is assigned for each correctly drawn symbol completed within the time limit. The total raw score is the number of correctly drawn symbols, with a maximum of 135. The raw scores were converted into age-scaled scores using the WAIS-IV manual.

*Verbal IQ* was assessed by means of the National Adult Reading Test (NART), based on Nelson and Willison (1991). The NART consists of 50 words with atypical phonemic pronunciation. Participants were instructed to slowly read aloud the list of words. Auditory recordings were made of the responses, which were individually rated by a native British speaker as either correct or incorrect according to the correct pronunciation as given by Google translate (2017, January 18). The NART error score consists of the total number of errors made on the complete NART. The Verbal IQ score that was used for analyses was calculated according to standard procedures: Estimated Verbal IQ = 129.0 −0.919 X NART error score.

### Markers of physical health

We assessed grip strength using a standard adjustable hand dynamometer (Takei Scientific Instruments). Standing in upright position, the participant was instructed to hold the dynamometer towards the ceiling with a completely outstretched arm, so that the shoulder and elbow were fully flexed at 180 degrees, hand palm facing the gaze direction. From this starting position, the participant was instructed to move their arm downwards in three seconds while squeezing the dynamometer with maximum force. A total of three measurements was recorded for the dominant and non-dominant hand, which was preceded by three practice trials for each hand. The highest value of the dominant hand was used for analyses. These raw scores were converted into standardised z-scores within age-and gender groups.

A physical activity questionnaire (New Zealand Physical Activities Survey Short Form; Sport and Recreation New Zealand, 2001) was included as a self-report measure of the participants’ habitual practice of physical activity. A composite score, calculated by adding the duration (in minutes) of moderate activity and two times the duration of vigorous activity, was used for analyses.

### Procedure

As mild hearing loss is a common condition in elderly people and the ability to clearly hear the stimuli is crucial for the aim of our study, the procedure started with a volume check. Participants listened to 20 randomly selected stimuli (10 real verbs and 10 pseudoverbs) through headphones and were asked to repeat what they heard. The experimenter paid special attention to correct pronunciation of the words’ suffices. Volume settings were adjusted if necessary.

Half of the participants started with the Pseudoverb block and the other half started with the Real Verb block. Instructions were identical in both blocks. After the participant read the instructions, the experimenter briefly summarized the procedure. Participants wore headphones and used the computer mouse to give their responses. Both blocks started with 33 practice trials, such that each possible word pair combination occurred three times. Participants received verbal feedback on their performance on the practice trials and only proceeded to the real experiment when they had a clear understanding of the task. The same procedure was repeated for the other block. Participants were instructed that the task in the second block was exactly the same as the previous one, only this time with real/pseudoverbs.

Each block took on average 30 minutes to complete, including the practice trials and two self-paced breaks. Participants were then tested on the additional measurements which were conducted in the following order: the Backward Digit Span Task; the Hand Grip Strength; the Physical Activity questionnaire; the Coding task and lastly the NART.

### Data analyses

The dependent variables are the accuracy and response time (RT) on the Correct syntax and Incorrect syntax trials^1^. The RT data for each participant in each condition was subjected to a ± 2 standard deviation trim, resulting in an exclusion of 5% of the data points in both groups. Lastly, one elderly participant was removed from further analyses due to excessively long RT’s (mean 2522, sd 1827, compared to the group mean 1164, sd 949)^2^. Only correct responses were included in the RT analyses. We analysed accuracy using a mixed-logit model in R (R Core Team, 2015), using the *Ime4* package (Bates, Mächler, Bolker & Walker, 2015). This method is most suitable for analysing categorical responses while excluding the necessity to conduct separate participant and item analyses (Jaeger, 2008). RT was analysed with a linear mixed model. The use of mixed effects models offers the opportunity to estimate effects and interactions of the experimental manipulations, or fixed effects, while simultaneously estimating parameters of the variance and covariance components of individual subjects and items as random effects (Kliegl, Wei, Dambacher, Yan, & Zhou, 2011).

To avoid multicollinearity in the regression models, we computed the Pearson’s correlation coefficients and p-values for our predictors using the corrplot package in R (Wei & Simko, 2016). Given that all correlations had a Spearman’s rank correlation coefficient <0.3, all predictors were included in the models.

The regression models for predicting both RT and Accuracy were based on the following predictors: Verb type (Pseudoverb and Real Verb); Syntax condition (Correct and Incorrect); Age group (younger and older); Working Memory; Processing Speed; Hand Grip; Physical Activity and Verbal IQ. Our categorical predictors Verb type, Syntax condition and Age group were all sum coded, such that the intercept of the model represents the grand mean (across all conditions) and the coefficients can directly be interpreted as main effects. Continuous variables were centred.

We began with a full model and then performed a step-wise “best-path” reduction procedure for the fixed effects to determine the simplest model that did not differ significantly from the full model in terms of variance explained (as described in Weatherholtz, Campbell-Kibler & Jaeger, 2014) using the dropl function from the stats package (version 3.4.2). We used a maximum random effects structure, allowing us to include intercepts for participants and items (“random intercepts”), as well as well as by-participants and by-item random slopes for the fixed effects. When the model did not converge with the fully expressed random effects structure, we simplified the random effects structure removing first the interactions, followed by the slopes which contributed least to the variance explained (Barr, Levy, Scheepers, & Tily, 2013).

Given that we were interested in the relationship between age and syntactic comprehension, the interactions that arose with the predictor Age group were further examined in post hoc analyses in which the regression models were applied to each Age group individually. Following this, the significant two way interactions in the post-hoc models were probed by testing each of the simple slopes for significance, using the *jtools* package in R (Long, 2018). Because the jtools package does not support lmer objects, we re-estimated the fixed effects using a Im function for our post hoc response time analyses.

## Results

### A. Group differences on individual differences measures

Table 2 provides an overview of the additional measurements for the younger and older Age group. In accordance with typical findings, the young participants outperform the older participants in Working Memory capacity and Processing Speed. To disentangle the effect of age from Processing Speed, the scaled scores were used in the analyses. However, for the sake of completeness, the raw scores are reported as well. The older participants performed significantly better in terms of Verbal IQ. There was no difference in Physical Activity or Hand Grip strength between both groups.

**Table 2.** Means and Standard Deviations of Additional measurements for the Young and Older Age group and the results of Comparisons between the Age groups (Independent Samples t-Test)

|  | Younger Age group<br>(N=50) |  | Older age group<br>(N= 50) |  | Comparison |  |
| --- | --- | --- | --- | --- | --- | --- |
|  | mean | sd | mean | sd | t | p |
| Working Memory | 5.57 | 1.57 | 4.57 | 1.59 | 3.1417 | 0.002 |
| Processing Speed | 80.34 | 14.91 | 62.92 | 16.48 | 5.5424 | < 0.001 |
| Processing Speed scaled | 11.52 | 2.91 | 12.52 | 16.48 | -1.7233 | 0.088 |
| Verbal IQ | 28.08 | 4.70 | 39.60 | 5.32 | -11.314 | < 0.001 |
| Physical Activity | 122.28 | 121.22 | 131.20 | 102.49 | -0.39736 | 0.692 |
| Hand grip | 25.92 | 7.54 | 26.99 | 8.85 | -0.65062 | 0.517 |

### B. Age differences in response accuracy for syntactic comprehension

We first discuss the main effect of Age group and Verb type on accuracy in order to answer our first two research questions concerning the effect of age on syntactic comprehension and the influence of semantic information. Following this, we will look at the effect of individual variation in our biomarkers on these results. Table 3 presents the results from the final mixed model predicting accuracy. This model was not significantly different from the full model (Full model = AIC: 6601.6, BIC 6915.4; Best model= AIC: 6598.8, BIC: 6897.7, p = 0.5447). Figure 2 (panel a) shows the group average of the proportion of correct responses given by the younger and the older age group for each of the two blocks. The younger age group obtained a mean accuracy of 95% (sd = 23) in the Real Verb block and a mean accuracy of 89% (sd =31) in the Pseudo Verb block. The older age group obtained a mean accuracy of 91% (sd = 29) and 89% (sd = 31) in the Real-and Pseudo Verb block respectively. The younger age group reached higher accuracy levels compared to the older age group in both the Real Verb and the Pseudoverb block (p < 0.001), suggesting that indeed there is age-related decline in syntactic comprehension accuracy. Generally, participants were less accurate in the Pseudoverb block compared to the Real Verb block (p = 0.001). The age-related decline in syntactic comprehension was stronger in the Real Verb block than the Pseudoverb block, as revealed by the significant Age group * Verb type interaction (p =0.039).

**Table 3.** Coefficient estimates, standard errors (SE), associated t values and significance levels for all predictors in the generalized mixed model predicting accuracy.

| Coefficient | Estimate | Std. Error | z value | p |  |
| --- | --- | --- | --- | --- | --- |
| **(Intercept)** | **3.072289** | **0.18477** | **16.628** | **< 0.001** | **\*\*\*** |
| **Working Memory** | **0.168661** | **0.063996** | **2.635** | **0.008** | **\*\*** |
| **Age group** | **1.576461** | **0.317789** | **4.961** | **< 0.001** | **\*\*\*** |
| **Verb type** | **-0.766071** | **0.233048** | **-3.287** | **0.001** | **\*\*** |
| **Syntax condition** | **0.289524** | **0.139877** | **2.07** | **0.038** | **\*** |
| Processing Speed | 0.025368 | 0.034454 | 0.736 | 0.462 |  |
| Handgrip | 0.034905 | 0.104121 | 0.335 | 0.737 |  |
| **Verbal IQ** | **0.113297** | **0.020378** | **5.56** | **< 0.001** | **\*\*\*** |
| Working Memory * Age group | -0.115867 | 0.128186 | -0.904 | 0.366 |  |
| Working Memory * Verb type | -0.015636 | 0.05394 | -0.29 | 0.772 |  |
| **Age group \* Verb type** | **0.577167** | **0.280153** | **2.06** | **0.039** | **\*** |
| Age group * Syntax condition | 0.263323 | 0.279732 | 0.941 | 0.347 |  |
| Verb type * Syntax condition | 0.08436 | 0.267458 | 0.315 | 0.752 |  |
| Working Memory * Syntax condition | 0.051873 | 0.052227 | 0.993 | 0.321 |  |
| Age group * Processing Speed | 0.007459 | 0.068913 | 0.108 | 0.914 |  |
| **Verb type \* Processing Speed** | **-0.065534** | **0.027935** | **-2.346** | **0.019** | **\*** |
| **Syntax condition \* Processing Speed** | **0.070299** | **0.027763** | **2.532** | **0.011** | **\*** |
| Age group * Handgrip | 0.200797 | 0.208306 | 0.964 | 0.335 |  |
| Verb type * Handgrip | -0.025933 | 0.078476 | -0.33 | 0.741 |  |
| Syntax condition * Handgrip | -0.001315 | 0.078096 | -0.017 | 0.987 |  |
| Age group * Verbal IQ | 0.028554 | 0.040691 | 0.702 | 0.483 |  |
| **Verb type \* Verbal IQ** | **0.098439** | **0.017262** | **5.703** | **< 0.001** | **\*\*\*** |
| Syntax condition * Verbal IQ | -0.009091 | 0.017171 | -0.529 | 0.596 |  |
| **Working Memory \* Age group \* Verb type** | **-0.278892** | **0.10793** | **-2.584** | **0.010** | **\*\*** |
| Age group * Verb type* Syntax condition | 0.410724 | 0.534602 | 0.768 | 0.442 |  |
| **Working Memory \* Age group \* Syntax condition** | **0.412985** | **0.104453** | **3.954** | **< 0.001** | **\*\*\*** |
| **Age group \* Verb type \* Processing Speed** | **0.210012** | **0.05587** | **3.759** | **< 0.001** | **\*\*\*** |
| Age group * Syntax condition * Processing Speed | -0.05964 | 0.05554 | -1.074 | 0.283 |  |
| **Verb type \* Syntax condition \* Processing Speed** | **-0.120479** | **0.054789** | **-2.199** | **0.029** | **\*** |
| Age group * Verb type * Handgrip | 0.233371 | 0.156948 | 1.487 | 0.137 |  |
| Age group * Syntax condition * Handgrip | -0.251511 | 0.156194 | -1.61 | 0.107 |  |
| **Verb type \* Syntax condition \* Handgrip** | **-0.602896** | **0.150283** | **-4.012** | **< 0.001** | **\*\*\*** |
| Age group * Verb type * Verbal IQ | -0.013328 | 0.034521 | -0.386 | 0.699 |  |
| Age group * Syntax condition * Verbal IQ | 0.037645 | 0.034338 | 1.096 | 0.273 |  |
| **Verb type \* Syntax condition \* Verbal IQ** | **0.136274** | **0.033519** | **4.066** | **< 0.001** | **\*\*\*** |
| Age group * Verb type * Syntax condition * Processing Speed | 0.194621 | 0.10959 | 1.776 | 0.076 | . |
| **Age group \* Verb type \* Syntax condition \* Handgrip** | **-0.685186** | **0.300617** | **-2.279** | **0.027** | **\*** |
| Age group * Verb type * Syntax condition * Verbal IQ | 0.118327 | 0.067046 | 1.765 | 0.078 | . |

*Signif. codes: 0 '\*\*\*' 0.001 '\*\*' 0.01 '\*' 0.05 '.' 0.1 ' ' 1* *Model includes a random intercept for items and subjects.*

**Figure 2.**
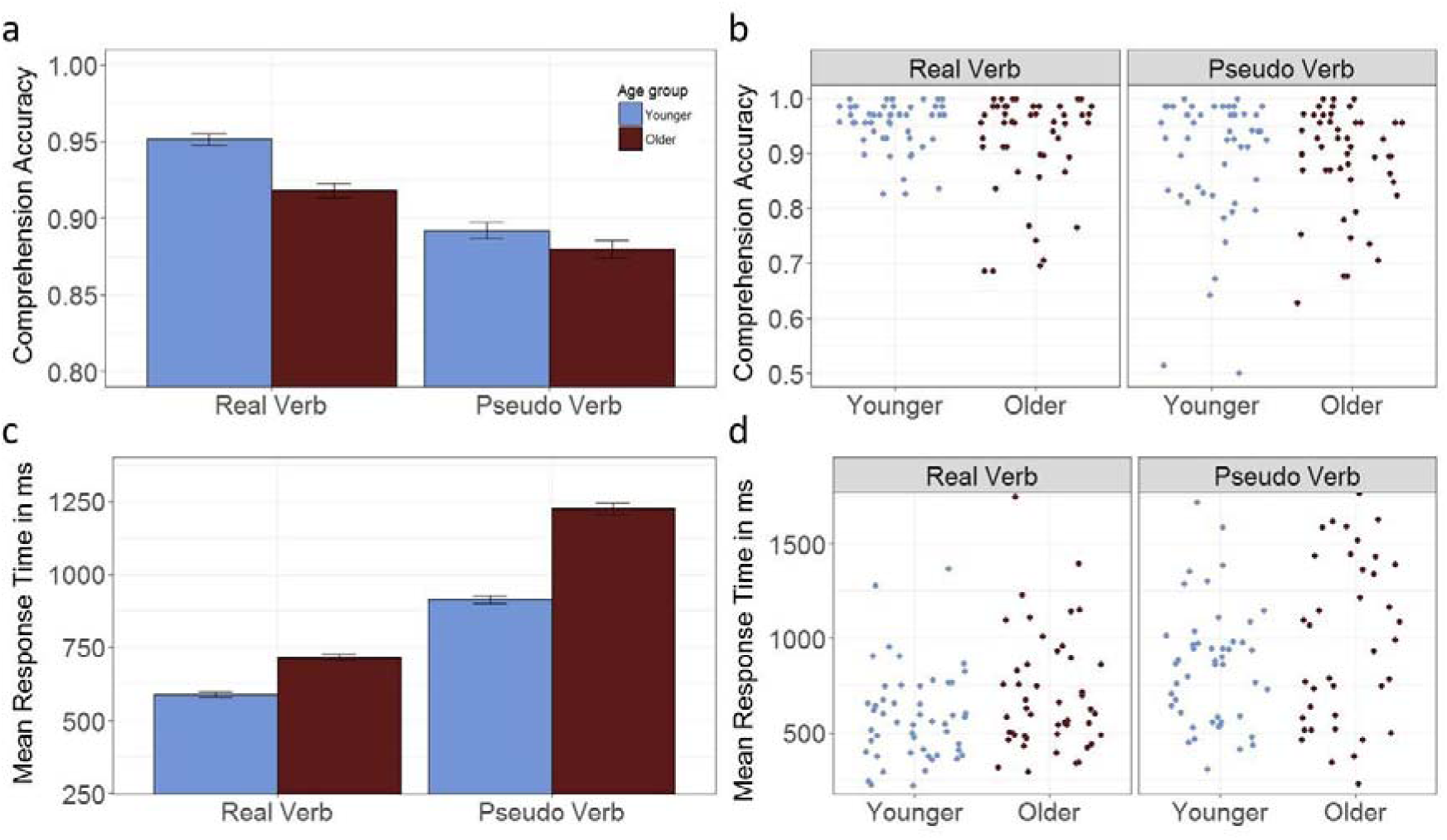
Age-related performance differences in accuracy (top row) and speed (bottom row) for syntactic comprehension. Group average proportion of correct comprehension per age group (a) and individual means b). Mean response times (RTs) to correct responses for the two age groups (c) and individual subjects (d). We have collapsed across the correct integration and incorrect integration condition in these graphs. Error bars are standard errors of the mean.

In addition to these group effects, there was individual variation in performance accuracy for both groups (shown in panel 2b)^3^. Of particular interest are interactions between individual difference measures and Age group, which were found for Processing speed and for Working memory. We turn to these next.

### Modulating effect of Processing Speed

There was a significant three-way interaction between Age group, Verb type and Processing Speed (p < 0.001), suggesting that processing speed modulates the effects of Age group and Verb Type on the accuracy of syntactic comprehension. To further examine this interaction, we ran a post hoc analysis in which the same model was applied to each age group individually. The results of this post hoc analysis are presented in Table 4. Linear regressions were created to visualise the interaction between Verb type and Processing Speed for each Age group separately. The left panel of Figure 3 shows the average accuracy as a function of Processing Speed in the younger age group for each Verb type separately. Accuracy was higher in the Real verb block compared to the Pseudoverb block. However, this effect of Verb type on accuracy did not depend on Processing Speed: there was no significant Verb type * Processing Speed interaction in the younger age group (p = 0.310). The right panel of Figure 3 shows the average accuracy as a function of Processing Speed for each of the two Verb types in the older age group. Similar to the younger age group, accuracy was higher in the Real verb block compared to the Pseudoverb block. However, the effect of Verb type on accuracy was qualified by an interaction between Verb type and Processing Speed in the older age group (p < 0.001). To determine whether this interaction was due to a larger influence of Processing Speed in the Real verb block relative to the Pseudoverb block, we ran a simple slope analysis for the influence of Processing Speed on accuracy for each level of Verb type (Real versus Pseudo). These post hoc z tests revealed the estimated beta coefficient in the Real verb block was significantly different from zero (B = 0.10; se = 0.06; z = −1.10, p = 0.08). In contrast, the beta coefficient in the Pseudoverb block was not significantly different from zero (B = −0.06; se = 0.06; z = −1.10; p = 0.27). Taken together, the results for older adults indicate that the effect of Processing Speed on accuracy is present in the Real verb block, but not in the Pseudoverb block. Older adults with higher Processing Speed performed better compared to older adults with lower Processing Speed in the Real Verb block. This suggests that higher Processing Speed in the older age group decreased the performance gap between younger and older participants in the Real Verb block. Note that we are using scaled Processing Speed scores so these effects cannot be attributed to effects of numerical age.

**Table 4A.** Coefficient estimates, standard errors, z values and p values of post hoc generalized mixed model predicting accuracy for the young age group. Model includes a random intercept for items and subjects.

| Coefficient | Estimate | Std. Error | z value | p |  |
| --- | --- | --- | --- | --- | --- |
| **(Intercept)** | **3.253857** | **0.170001** | **19.14** | **< 0.001** | **\*\*\*** |
| Working Memory | 0.115142 | 0.080689 | 1.427 | <0.156 |  |
| **Verb type** | **-1.19055** | **0.262885** | **-4.529** | **< 0.001** | **\*\*\*** |
| **Syntax condition** | **0.48613** | **0.123556** | **3.934** | **< 0.001** | **\*\*\*** |
| Processing Speed | 0.028889 | 0.042403 | 0.681 | 0.496 |  |
| Handgrip | 0.139061 | 0.121419 | 1.145 | 0.252 |  |
| **Verbal IQ** | **0.129433** | **0.028112** | **4.604** | **< 0.001** | **\*\*\*** |
| **Working Memory \* Verb type** | **-0.15881** | **0.072449** | **-2.192** | **0.028** | **\*** |
| **Verb type \* Syntax condition** | **-0.85685** | **0.248179** | **-3.453** | **< 0.001** | **\*\*\*** |
| **Working Memory \* Syntax condition** | **0.263092** | **0.068853** | **3.821** | **< 0.001** | **\*\*\*** |
| Verb type * Processing Speed | 0.041828 | 0.041219 | 1.015 | 0.310 |  |
| Syntax condition * Processing Speed | 0.039648 | 0.04119 | 0.963 | 0.336 |  |
| Verb type * Handgrip | 0.082392 | 0.108652 | 0.758 | 0.448 |  |
| Syntax condition * Handgrip | -0.12103 | 0.108464 | -1.116 | 0.264 |  |
| **Verb type \* Verbal IQ** | **0.088525** | **0.027773** | **3.187** | **0.001** | **\*\*** |
| Syntax condition * Verbal IQ | 0.009718 | 0.027694 | 0.351 | 0.726 |  |
| Verb type * Syntax condition * Processing Speed | -0.02185 | 0.080702 | -0.271 | 0.787 |  |
| **Verb type \* Syntax condition \* Handgrip** | **-0.97036** | **0.219806** | **-4.415** | **< 0.001** | **\*\*\*** |
| **Verb type \* Syntax condition \* Verbal IQ** | **0.199873** | **0.053268** | **3.752** | **< 0.001** | **\*\*\*** |
| **4B. Coefficient estimates, standard errors, z values and p values of post hoc generalized mixed model predicting accuracy for the older age group. Model includes a random intercept for items and subjects.** |  |  |  |  |  |
| Coefficients | Estimate | Std. Error | z value | p |  |
| **(Intercept)** | **2.72263** | **0.15951** | **17.069** | **< 0.001** | **\*\*\*** |
| **Working Memory** | **0.23384** | **0.10064** | **2.324** | **0.020** | **\*** |
| **Verb type** | **-0.49327** | **0.18791** | **-2.625** | **0.009** | **\*\*** |
| Syntax condition | 0.11606 | 0.09853 | 1.178 | 0.239 |  |
| Processing Speed | 0.02056 | 0.05528 | 0.372 | 0.710 |  |
| Handgrip | -0.07129 | 0.17354 | -0.411 | 0.681 |  |
| **Verbal IQ** | **0.09965** | **0.02942** | **3.388** | **< 0.001** | **\*\*\*** |
| Working Memory * Verb type | 0.12121 | 0.08021 | 1.511 | 0.131 |  |
| Verb type * Syntax condition | 0.2408 | 0.19381 | 1.242 | 0.214 |  |
| Working Memory * Syntax condition | -0.14907 | 0.0784 | -1.901 | 0.057 | . |
| **Verb type \* Processing Speed** | **-0.16688** | **0.03747** | **-4.454** | **< 0.001** | **\*\*\*** |
| **Syntax condition \* Processing Speed** | **0.0971** | **0.037** | **2.624** | **0.009** | **\*\*** |
| Verb type * Handgrip | -0.13955 | 0.11276 | -1.238 | 0.216 |  |
| Syntax condition * Handgrip | 0.1227 | 0.112 | 1.096 | 0.273 |  |
| **Verb type \* Verbal IQ** | **0.10528** | **0.02068** | **5.09** | **< 0.001** | **\*\*\*** |
| Syntax condition * Verbal IQ | -0.0271 | 0.02038 | -1.33 | 0.184 |  |
| **Verb type \* Syntax condition \* Processing Speed** | **-0.20898** | **0.07378** | **-2.832** | **0.005** | **\*\*** |
| Verb type * Syntax condition * Handgrip | -0.26974 | 0.20453 | -1.319 | 0.187 |  |
| Verb type * Syntax condition * Verbal IQ | 0.07653 | 0.04083 | 1.874 | 0.061 | . |
| *Signif. codes: 0 '\*\*\*' 0.001 '\*\*' 0.01 '\*' 0.05 '.' 0.1 ' ' 1* |  |  |  |  |  |

**Figure 3.**
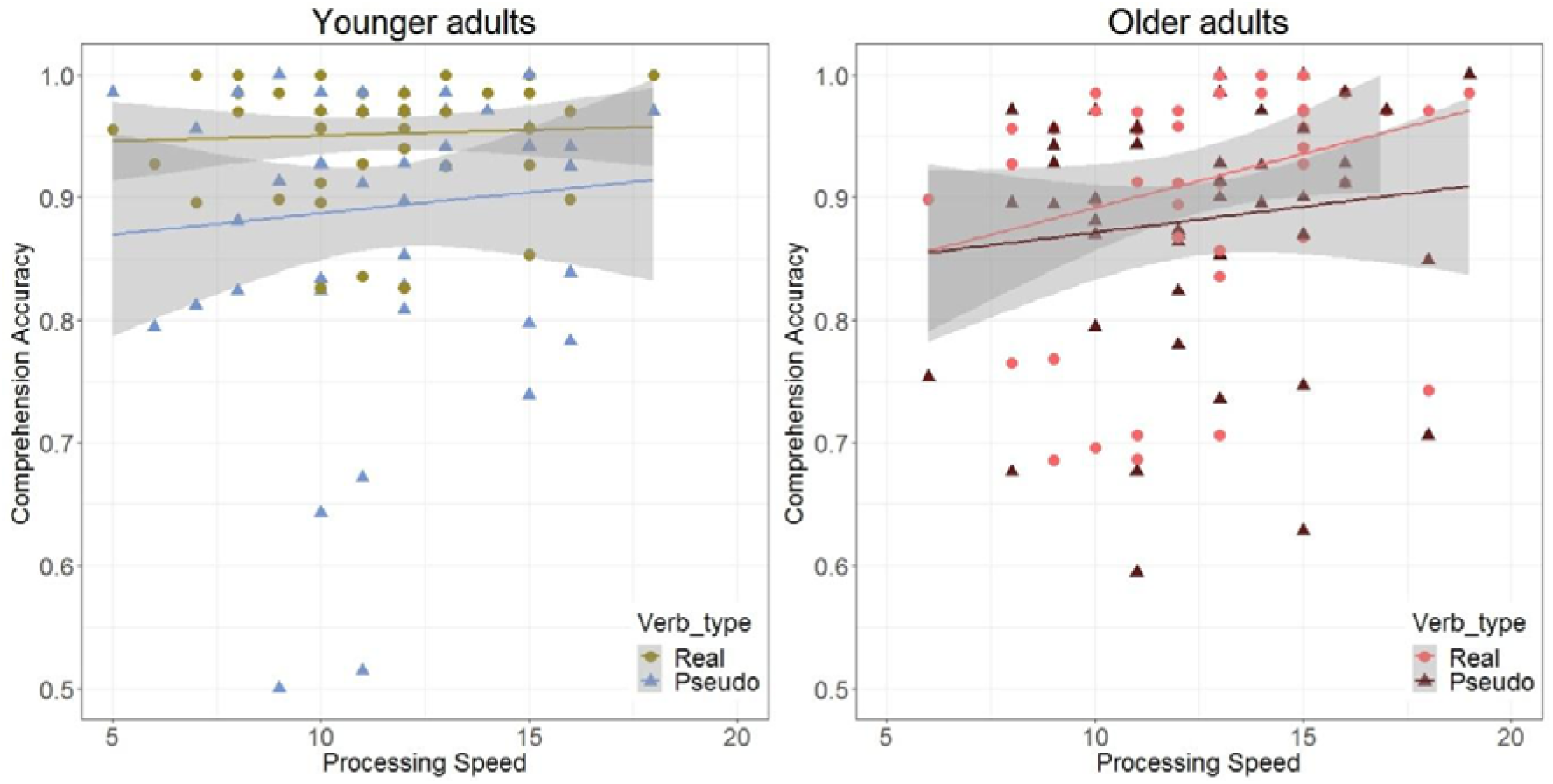
Processing speed modulates syntactic comprehension accuracy in the older Age group. Three-way interaction between Age group, Verb type and Processing Speed depicted through a linear regression with accuracy as predicted by Processing Speed in the Real verb and Pseudoverb block for each age group separately. The left panel shows the younger age group, the right panel shows the older age group. Processing speed influenced the effect of Verb type on accuracy in the older age group, but not in the younger age group.

### Modulating effect of Working Memory

To assess whether Working Memory modulates the effect of Age group on accuracy, we looked at interactions between Working Memory and Age group. There was a significant three-way interaction between Age group, Working Memory and Verb type (p = 0.010), which was further examined in a post hoc analysis by applying the same model to each age group individually (see Table 4). The left panel of Figure 4 shows the linear regressions of Working Memory predicting accuracy for the two different Verb types in the younger age group. The effect of Verb type on accuracy was influenced by Working Memory, as evidenced by the significant Working Memory * Verb type interaction (p = 0.028). To further interpret this interaction, we performed a simple slopes analysis for the effect of Working Memory in each of the two Verb types. In the Real verb block the estimated beta coefficient was significantly different from zero (B = 0.19; se = 0.09; z = 2.08; p = 0.04). In contrast, in the Pseudoverb block the beta coefficient was not significantly different from zero (B = 0.04; se = 0.08; z = 0.43; p = 0.67). This suggests that the effect of Working Memory on accuracy was only present in the Real verb block, such that younger adults with higher Working Memory scores obtained a higher accuracy in the Real verb block compared to younger adults with lower Working Memory scores. The right panel of Figure 4 shows the linear regressions of Working Memory predicting accuracy for the two different Verb types in the older age group. Working Memory influenced accuracy in the older age group (p = 0.020), such that older adults with higher Working Memory scores performed better than older adults with lower Working Memory scores. However, the effect of Working Memory did not differ across Verb type: there was no significant Working Memory * Verb type interaction (p = 0.131).

**Figure 4.**
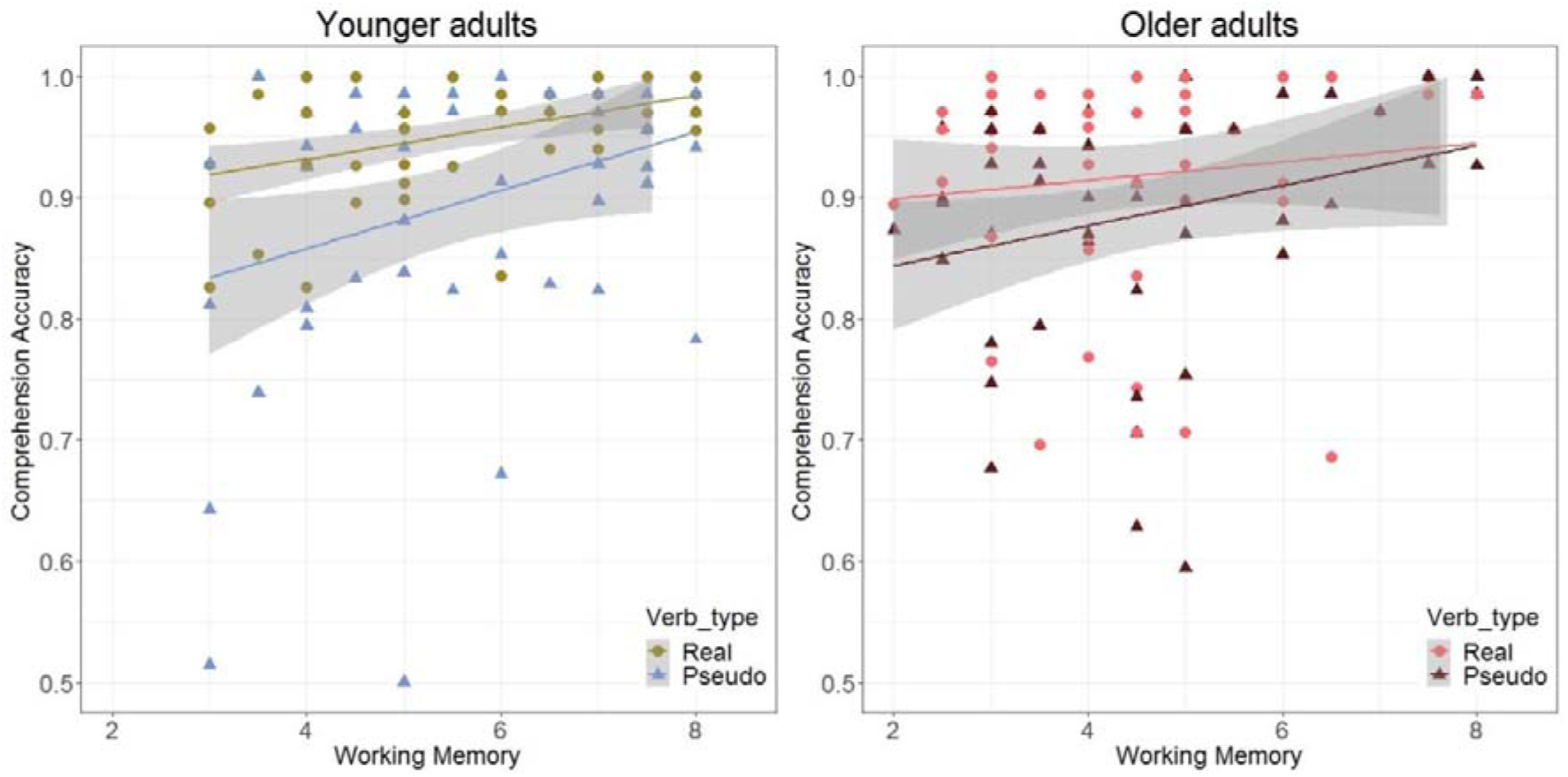
Working Memory differentially effects syntactic comprehension accuracy depending on Age group. The three-way interaction between Age group, Verb type and Working Memory, depicted by a linear regression between accuracy and Working Memory grouped by Verb type in the younger age group (left panel) and the older age group (right panel). Lower Working Memory in the young adults was associated with decreased accuracy in the Real verb block. The relationship between Working Memory and accuracy was not different for the two Verb types in the older age group.

Notably, there was an additional significant three-way interaction between Age group, Working Memory and Syntax condition (p < 0.001), which was driven by a significant interaction between Working Memory and Syntax condition in the younger age group (p < 0.001), but not in the older age group (p = 0.057). The post hoc simple slopes analyses revealed a non-significant effect of Working Memory on accuracy in the Correct Syntax condition (B = −0.02; se = 0.08; z = −0.19; p = 0.85) and a significant effect of Working Memory in the Incorrect Syntax condition (B = 0.25; se = 0.09; z = 2.72; p = 0.01). These results indicate that lower Working Memory was associated with lower task performance in the Incorrect Syntax condition in the younger age group.

Overall, this suggests that in younger adults, a lower working memory span is associated with a relative disadvantage in performance in comprehending real verb phrases and in correctly identifying morpho-syntactically incorrect phrases. In contrast, higher Working Memory was associated with higher accuracy in the older age group regardless of Verb type or Syntax condition.

### Modulating effect of Handgrip strength

We found a significant four way interaction between Age group, Verb type, Syntax condition and Handgrip (p = 0.027). Post hoc analyses revealed this effect was driven by a significant interaction between Verb type, Syntax condition and Handgrip in the young age group (p < 0.001). There was no significant interaction between Verb type, Syntax condition and Handgrip in the older age group (p= 0.187). In the younger age group, accuracy in the Incorrect Syntax condition of the Pseudoverb block was particularly low and modulated by variability in handgrip scores.

### C. Age differences in response time for syntactic comprehension

Similar to the accuracy results, we will first discuss the overall group differences in response time in relation to Verb type before we discuss how these group differences can be further explained by the inter individual variability markers. Table 5 presents the results of the best linear mixed model predicting response times. This model was not significantly different from the full model (Full model = AIC: 183053 BIC 183510; Best model= AIC: 183034 BIC: 183395, p = 0.902). Figure 2 (panel c) shows the mean response times in ms on the Pseudoverb and Real verb block for both age groups. The mean response time in the younger age group was 757 ms (sd = 529) in the Real Verb block and 979 ms (sd = 731) in the Pseudoverb block. In the older age group, the mean response time was 871 ms (sd = 695) in the Real Verb block and 1270 ms (sd = 982) in the Pseudoverb block. The older age group took longer to respond than the younger age group (p < 0.001). In addition, response times were significantly longer in the Pseudoverb block compared to the Real verb block (p < 0.001). Age-related decline in response times was larger for the Pseudoverb block compared to the Real verb block, as revealed by the Age group * Verb type interaction (p = 0.008). Post hoc analyses within each age group revealed that the effect of Verb type exists in both age groups (see Table 6). However, as can be seen in Figure 2c, the effect is larger in the older age group.

**Table 5.**
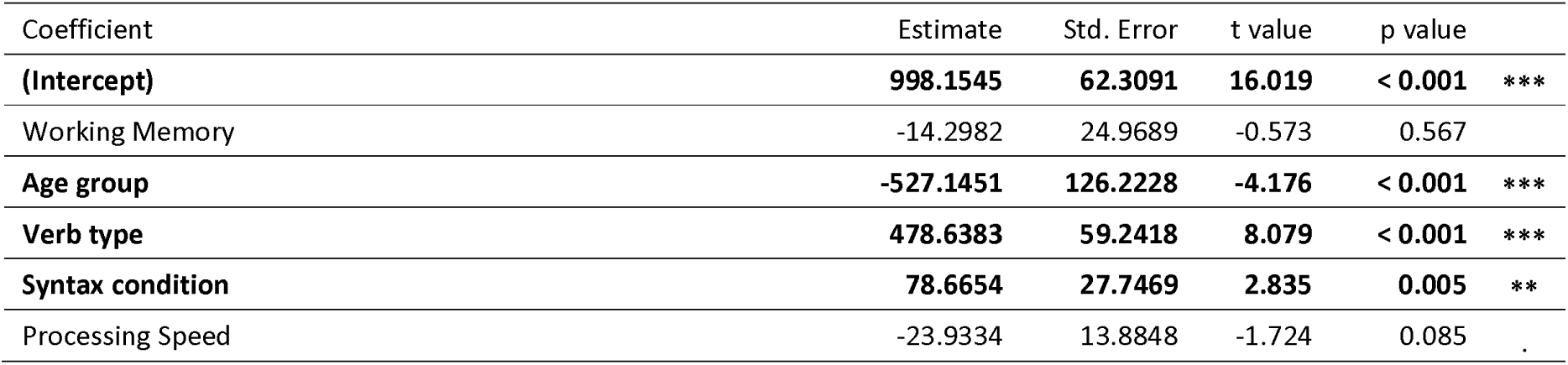
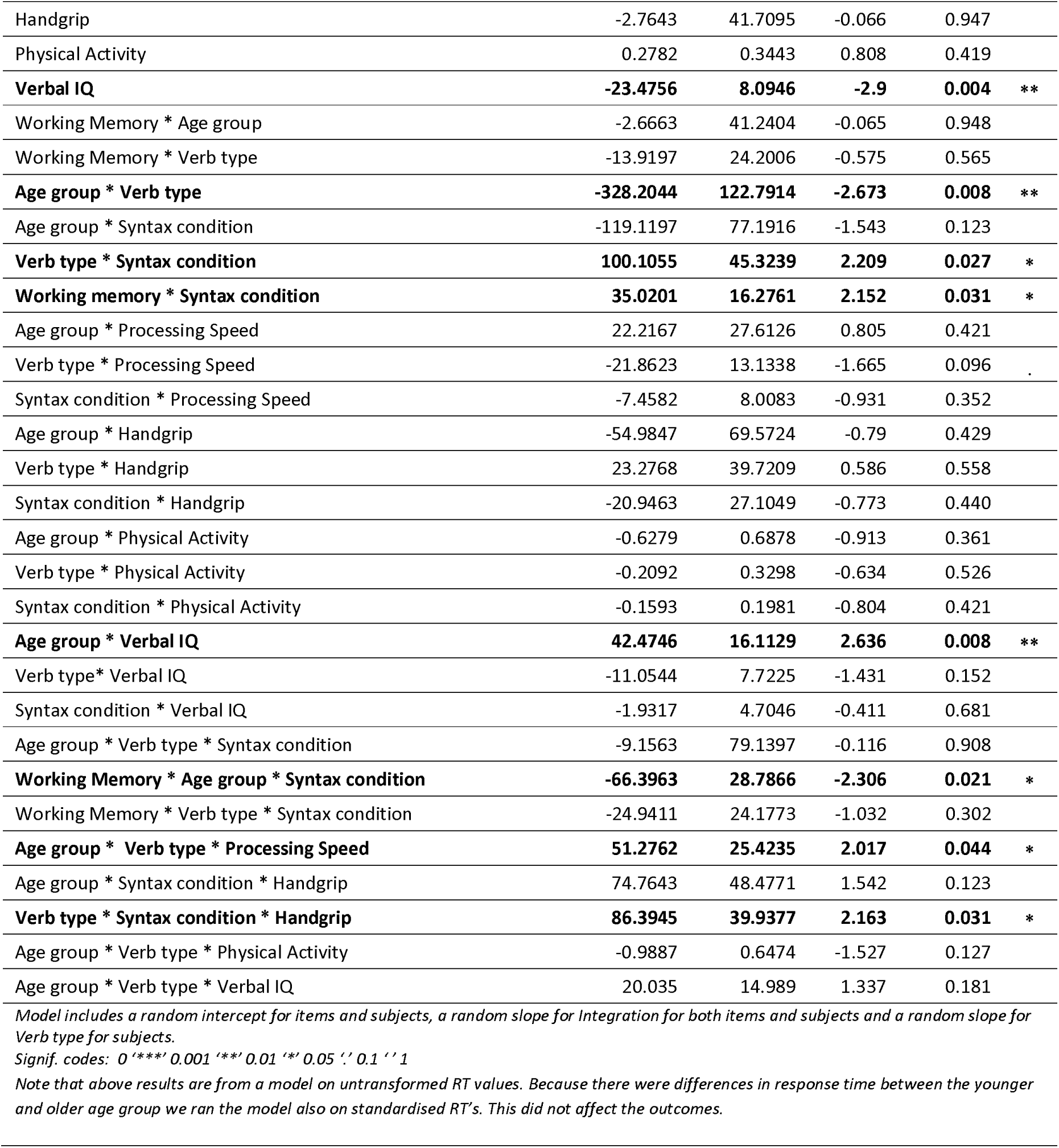
Coefficient estimates, standard errors (SE), associated t values and p values for all predictors of linear mixed model predicting response time.

**Table 6.A.**
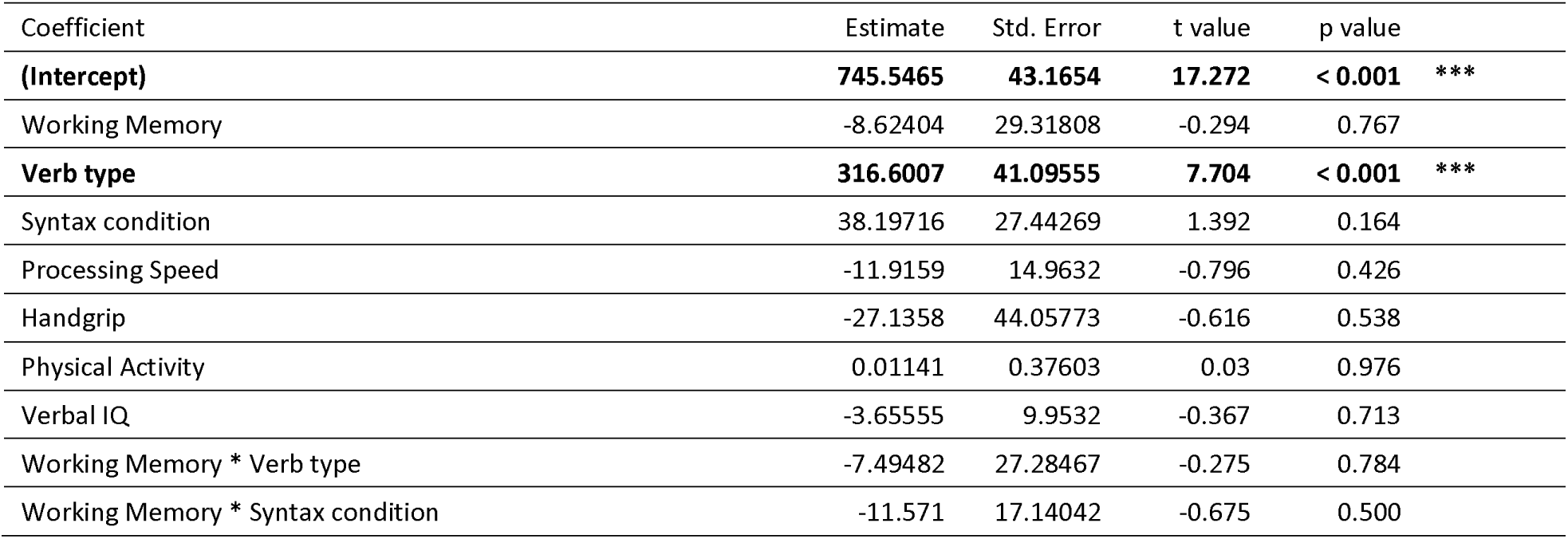
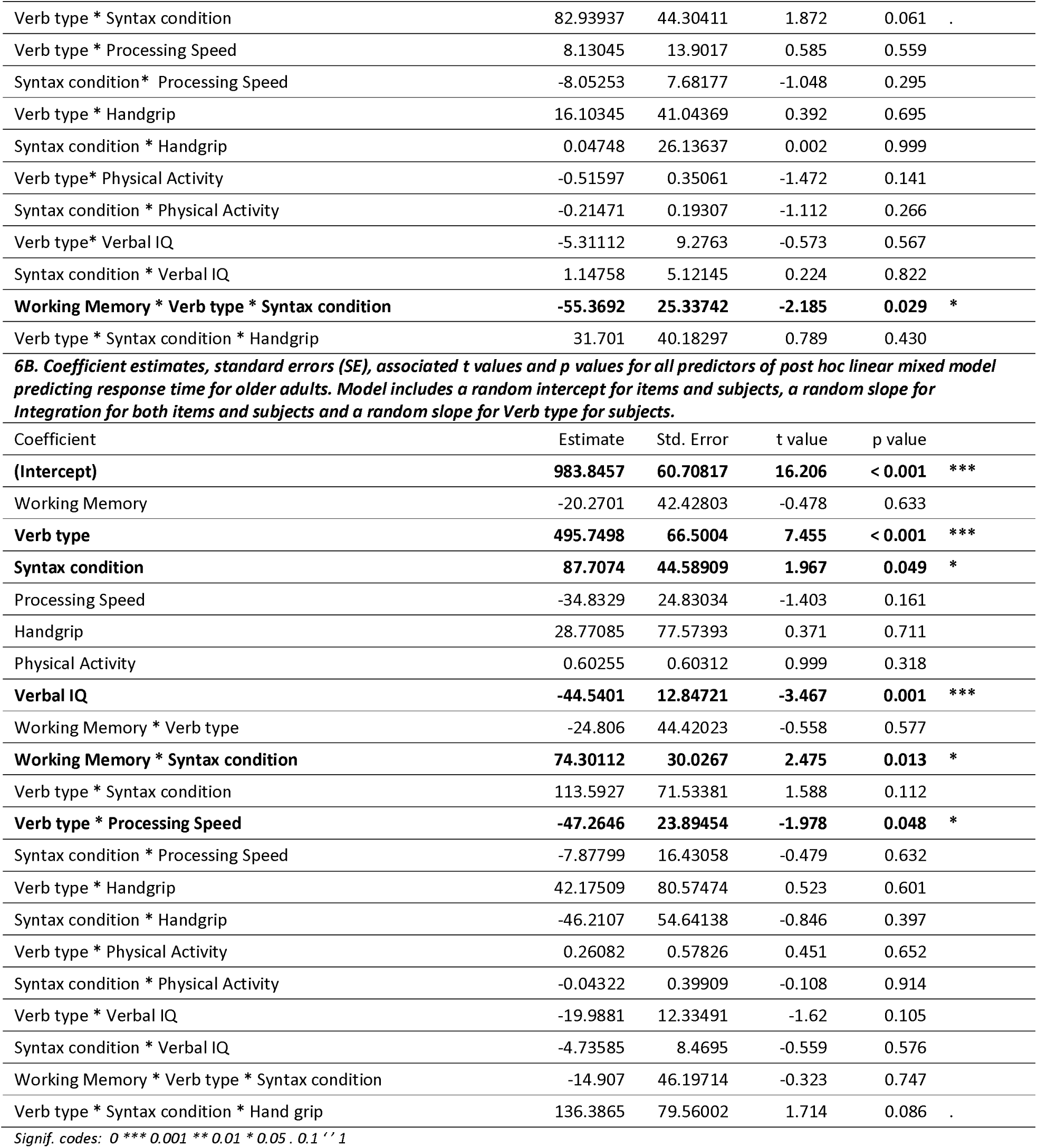
Coefficient estimates, standard errors (SE), associated t values and p values for all predictors of post hoc linear mixed model predicting response time for young adults. Model includes a random intercept for items and subjects, a random slope for Integration for both items and subjects and a random slope for Verb type for subjects.

| Coefficient | Estimate | Std. Error | t value | p value |
| --- | --- | --- | --- | --- |
| **(Intercept)** | **745.5465** | **43.1654** | **17.272** | **< 0.001 \*\*\*** |
| Working Memory | -8.62404 | 29.31808 | -0.294 | 0.767 |
| **Verb type** | **316.6007** | **41.09555** | **7.704** | **< 0.001 \*\*\*** |
| Syntax condition | 38.19716 | 27.44269 | 1.392 | 0.164 |
| Processing Speed | -11.9159 | 14.9632 | -0.796 | 0.426 |
| Handgrip | -27.1358 | 44.05773 | -0.616 | 0.538 |
| Physical Activity | 0.01141 | 0.37603 | 0.03 | 0.976 |
| Verbal IQ | -3.65555 | 9.9532 | -0.367 | 0.713 |
| Working Memory * Verb type | -7.49482 | 27.28467 | -0.275 | 0.784 |
| Working Memory * Syntax condition | -11.571 | 17.14042 | -0.675 | 0.500 |

|  |  |  |  |  |  |
| --- | --- | --- | --- | --- | --- |
| Verb type * Syntax condition | 82.93937 | 44.30411 | 1.872 | 0.061 | . |
| Verb type * Processing Speed | 8.13045 | 13.9017 | 0.585 | 0.559 |  |
| Syntax condition * Processing Speed | -8.05253 | 7.68177 | -1.048 | 0.295 |  |
| Verb type * Handgrip | 16.10345 | 41.04369 | 0.392 | 0.695 |  |
| Syntax condition * Handgrip | 0.04748 | 26.13637 | 0.002 | 0.999 |  |
| Verb type* Physical Activity | -0.51597 | 0.35061 | -1.472 | 0.141 |  |
| Syntax condition * Physical Activity | -0.21471 | 0.19307 | -1.112 | 0.266 |  |
| Verb type* Verbal IQ | -5.31112 | 9.2763 | -0.573 | 0.567 |  |
| Syntax condition * Verbal IQ | 1.14758 | 5.12145 | 0.224 | 0.822 |  |
| **Working Memory \* Verb type \* Syntax condition** | **-55.3692** | **25.33742** | **-2.185** | **0.029** | **\*** |
| Verb type * Syntax condition * Handgrip | 31.701 | 40.18297 | 0.789 | 0.430 |  |
| **6B. Coefficient estimates, standard errors (SE), associated t values and p values for all predictors of post hoc linear mixed model predicting response time for older adults. Model includes a random intercept for items and subjects, a random slope for Integration for both items and subjects and a random slope for Verb type for subjects.** |  |  |  |  |  |
| Coefficient | Estimate | Std. Error | t value | p value |  |
| **(Intercept)** | **983.8457** | **60.70817** | **16.206** | **< 0.001** | **\*\*\*** |
| Working Memory | -20.2701 | 42.42803 | -0.478 | 0.633 |  |
| **Verb type** | **495.7498** | **66.5004** | **7.455** | **< 0.001** | **\*\*\*** |
| **Syntax condition** | **87.7074** | **44.58909** | **1.967** | **0.049** | **\*** |
| Processing Speed | -34.8329 | 24.83034 | -1.403 | 0.161 |  |
| Handgrip | 28.77085 | 77.57393 | 0.371 | 0.711 |  |
| Physical Activity | 0.60255 | 0.60312 | 0.999 | 0.318 |  |
| **Verbal IQ** | **-44.5401** | **12.84721** | **-3.467** | **0.001** | **\*\*\*** |
| Working Memory * Verb type | -24.806 | 44.42023 | -0.558 | 0.577 |  |
| **Working Memory \* Syntax condition** | **74.30112** | **30.0267** | **2.475** | **0.013** | **\*** |
| Verb type * Syntax condition | 113.5927 | 71.53381 | 1.588 | 0.112 |  |
| **Verb type \* Processing Speed** | **-47.2646** | **23.89454** | **-1.978** | **0.048** | **\*** |
| Syntax condition * Processing Speed | -7.87799 | 16.43058 | -0.479 | 0.632 |  |
| Verb type * Handgrip | 42.17509 | 80.57474 | 0.523 | 0.601 |  |
| Syntax condition * Handgrip | -46.2107 | 54.64138 | -0.846 | 0.397 |  |
| Verb type * Physical Activity | 0.26082 | 0.57826 | 0.451 | 0.652 |  |
| Syntax condition * Physical Activity | -0.04322 | 0.39909 | -0.108 | 0.914 |  |
| Verb type * Verbal IQ | -19.9881 | 12.33491 | -1.62 | 0.105 |  |
| Syntax condition * Verbal IQ | -4.73585 | 8.4695 | -0.559 | 0.576 |  |
| Working Memory * Verb type * Syntax condition | -14.907 | 46.19714 | -0.323 | 0.747 |  |
| Verb type * Syntax condition * Hand grip | 136.3865 | 79.56002 | 1.714 | 0.086 | . |
| Signif. codes: 0 '***' 0.001 '**' 0.01 '*' 0.05 '.' 0.1 ' ' 1 |  |  |  |  |  |

In addition to these group effects, we were interested in the moderating influence of our cognitive and physical markers, to further explain the individual variation in reaction times that was present in both groups (shown in panel 2d)^4^. Of particular interests are interactions that modulate the effect of Age group on response time, which were found for Processing Speed and Working Memory. We turn to a description of these results next.

### Modulating effect of Processing Speed

To assess whether the effect of Processing Speed on response time was different for younger and older adults, we looked at interactions between Age group and Processing Speed. Similar to our accuracy analyses, we found an interaction between Age group, Verb type and Processing Speed (p = 0.044). To investigate the nature of this interaction, we ran a post hoc analysis in which the model predicting response times was applied to each age group individually. The results of this post hoc analysis are presented in Table 6. The left panel of Figure 5 shows that in the younger age group, response times were shorter in the Real verb block compared to the Pseudoverb block. This effect of Verb type on response time did not depend on Processing Speed: there was no significant interaction between Processing Speed and Verb type in the younger age group (p = 0.559). In the older age group (right panel of Figure 5), response times were shorter in the Real verb block compared to the Pseudoverb block. However, the effect of Verb type on response times was moderated by Processing Speed: there was a significant Verb type * Processing Speed interaction in the older age group (p= 0.048). To investigate this interaction, we tested the slope for the effect of Processing Speed on response time for each Verb type separately. These post hoc t tests revealed the estimated beta coefficient in the Real verb block was not significantly different from zero (B = −8.73; se = 6; t = −1.46; p = 0.15). In contrast, the beta coefficient in the Pseudoverb block was significantly different from zero (B = −57.89; se = 6.15; t = −9.42; p < 0.001). This suggests that the relative increase in response time in the Pseudoverb block was elevated in older adults with lower Processing Speed.

**Figure 5.**
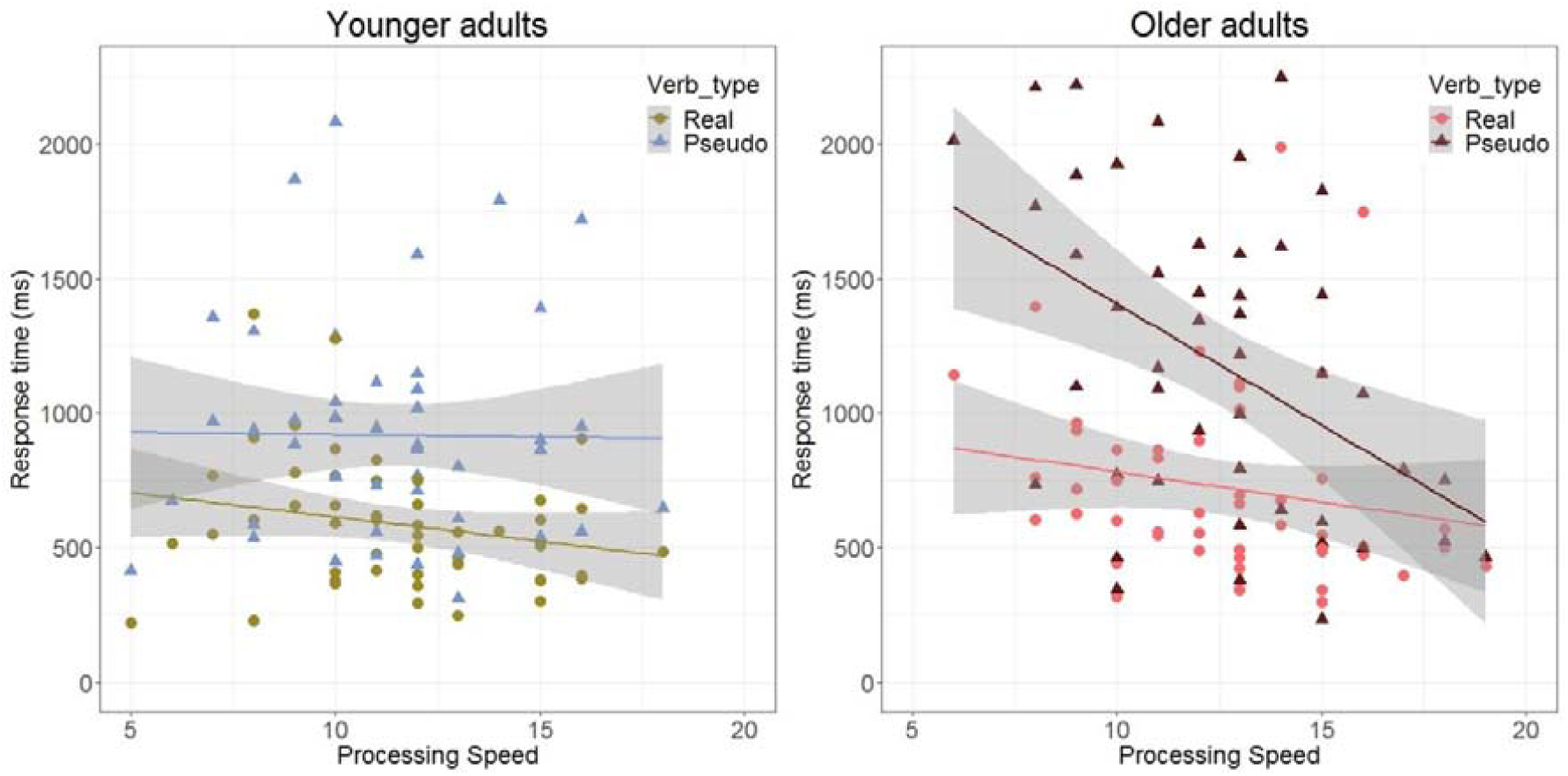
Processing Speed differentially effects response time depending on age group. The three-way interaction between Age group, Verb type and Processing Speed, depicted through a linear regression with response time as predicted by Processing Speed in the Real verb and Pseudoverb block for each age group separately. The left panel shows the younger age group, the right panel shows the older age group. In the younger age group, the effect of Verb type on response time was not influenced by Processing Speed. In contrast, the effect of Verb type on response time was different at different levels of Processing Speed in the older age group.

### Moderating effect of Working Memory

To investigate whether Working Memory differentially affects response times in younger and older individuals, we looked at interactions between Working Memory and Age group. There was a significant interaction between Age group, Working Memory and Syntax condition (p = 0.021). As can be seen in the left panel of Figure 6, the response times in the younger age group did not differ across conditions and Working Memory did not influence the response times: there was no significant interaction between Working Memory and Syntax condition in the younger age group (p = 0.5; see Table 6). As can be seen in the right panel of Figure 6, the effect of Syntax Condition was moderated by Working Memory in the older age group. Specifically, response times were shorter in the Correct Syntax condition compared to the Incorrect Syntax condition, but this difference is driven by older adults with higher working memory: there was a significant interaction between Working Memory and Syntax condition (p = 0.013). To determine whether the effect of Syntax condition was larger in the Correct Syntax condition relative to the Incorrect Syntax condition, we tested the simple slopes of the influence of Working Memory in each Syntax condition against zero. The post hoc t tests revealed the simple slope in the Correct syntax condition was significantly different from zero (B = −58.40; se = 10.12; t = −5.77; p < 0.001). In contrast, the simple slope in the Incorrect Syntax condition was not significantly different from zero (B = 9.51; se = 10.02; t = 0.95; p = 0.34). Overall, this suggests that for older adults, higher working memory was associated with slower response times in the Correct Syntax condition, while for younger adults, working memory did not influence the response times.

**Figure 6.**
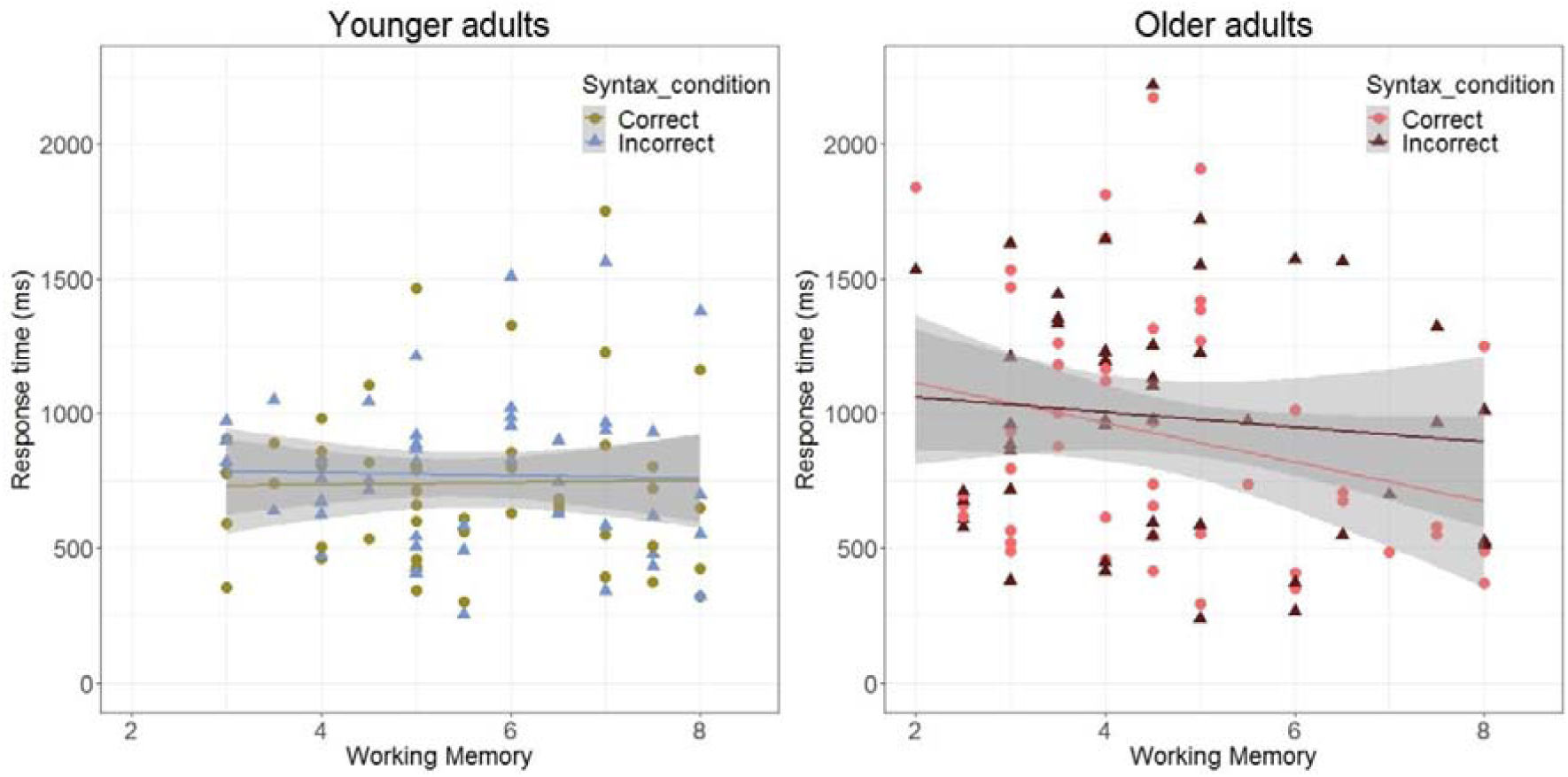
Working Memory differentially effects response time depending on age group. The three-way interaction between Age group, Syntax condition and Working Memory, depicted by a linear regression between response time and Working Memory grouped by Syntax condition in the younger age group (left panel) and the older age group (right panel). Working Memory did not differentially affect response times depending on Syntax condition in the younger age group. In the older age group, there was a significant decrease in response time in older adults with high working memory in the Correct Syntax condition.

### Moderating effect of Verbal IQ

We found an interaction between Age Group and Verbal IQ (p = 0.008), such that a higher Verbal IQ score was associated with faster response times for older adults, but not for young adults.

## Discussion

Our study was designed to investigate whether there is decline in syntactic comprehension in healthy ageing. We investigated elementary syntactic comprehension of phrases such as “I cook” and “I spuff”. We demonstrated the following three key findings: 1) there is decline in syntactic comprehension of healthy older adults compared to young adults, in accuracy as well as response times; 2) the age-related decline in the accuracy of syntactic comprehension is stronger for phrases with real verbs, while the age-related decline in the response times of syntactic comprehension is stronger for phrases with pseudoverbs; 3) there is a high degree of individual variation in age-related decline, which is explained in part by differences in working memory and processing speed.

The modulations of processing speed and working memory on syntactic comprehension present a complex picture, which can be summarized as follows. In young adults, performance was not affected by processing speed. This was true for accuracy as well as response time. In older adults, processing speed influenced syntactic comprehension, both in terms of accuracy and response time. However, processing speed differentially influences performance on accuracy and response time depending on the level of lexical-semantic information provided. Specifically, in real verb sentences, processing speed aids accuracy of syntactic judgements, whereas in pseudoverb sentences, processing speed aids response times. The moderating influence of working memory on comprehension performance was different for the two age groups as well. In older adults, working memory aids accuracy, an advantage which was not dependent on the level of lexical-semantic information provided (whereas for young adults it was). Moreover, working memory aids response times in syntactically correct sentences. We discuss these effects below in the context of our key findings.

We have convincingly demonstrated that there is age-related decline in syntactic comprehension when processing two-word phrases with real verbs in our syntactic comprehension experiment. The effects were demonstrated in accuracy as well as response times: older adults were less accurate and slower than young adults. Previous literature on syntactic comprehension in older adults has predominantly used semantically meaningful sentences with complex syntactic structures. Most of these studies did not show age-related decline in processing these sentences (Campbell et al., 2016; Davis et al., 2014; Meunier et al., 2014; Samu et al., 2017; Shafto & Tyler, 2014b; Shafto et al., 2014), although some studies did (Antonenko et al., 2013; Obler, Fein, Nicholas, & Albert, 1991). Our results are in line with the latter set of studies. A new element in the results of the current study is that age related decline in syntactic comprehension was demonstrated in a context where complexity was reduced to the bare minimum: syntactic agreement of pronoun and verb.

A possible explanation for the divergence in results of the current study compared to many previous findings of preserved syntactic comprehension is that the measure of syntactic comprehension used in the current study may draw on a different aspect of syntax. Studies that capitalize on syntactic ambiguity evaluate comprehension by asking questions about the thematic roles assigned to the agent or patient in the sentence (i.e ‘who is doing what’, e.g ‘what is the gender of the agent in the sentence’). A correct response requires comprehension of the full sentence structure, which indirectly requires comprehension of the syntactic structure. In contrast, the measure of syntactic comprehension in the current study focuses on evaluating syntactic agreement. This study thus taps into a different aspect of syntactic processing: grammaticality judgements for minimal phrases with and without meaning. Specifically, in the context of Friederici’s (2000) neurocognitive model of auditory sentence processing, the current study arguably taps into the initial phases of sentence processing of local syntactic structure building and thematic role assignment based on morpho-syntactic information indicating agreement between different elements within a phrase. In contrast, syntactic ambiguity paradigms (as used by Campbell et al., 2016; Davis, Zhuang, Wright, & Tyler, 2014; Meunier, Stamatakis, & Tyler, 2014; Samu et al., 2017; Shafto & Tyler, 2014) tap into later processes of syntactic revision and late integration (although see Antonenko et al., 2013 for a study with a syntactic ambiguity paradigm that did find age-related decline). Different aspects of syntactic processing do not necessarily undergo a similar trajectory of change over the course of aging. The current study only enables us to draw conclusions on those aspects of syntax that were manipulated in our experiment design. Moreover, our task is a metalinguistic task that requires post-interpretive processing. For a review on the possible effects of ageing on the added processes involved in post-interpretive tasks, please see a review by Peelle (in press).

Our second key finding is that the pattern and extent of age-related decline is influenced by the level of lexical semantic information provided. The reduction of lexical semantic content by using pseudoverbs instead of real verbs increased the difficulty of the task, as evidenced by the reduced accuracy and increased response times in both age groups. Older adults were slower and less accurate in comprehending both real verb and pseudoverb phrases. In terms of accuracy, this relative performance drop was largest in the real verb phrases compared to the pseudoverb phrases. In terms of response time, the age-related decline was largest in the pseudoverb phrases compared to the real verb phrases. Older and younger adults likely used a different strategy: while younger adults more often adopt a strategy that emphasizes speed, older adults tend to act more error aversive than younger adults (de Jong et al., 2018). Indeed, it has been suggested previously that older adults prioritize accurate responses over fast responses (Forstmann et al., 2011; Starns & Ratcliff, 2010).

One possible interpretation of this pattern of findings is that decline in syntactic comprehension is strongest in the absence of lexical-semantic information, which causes older adults to produce slower responses in order to make more accurate decisions. This interpretation of the results could shed some light on why some previous studies did not show any decline in syntactic processing when syntactic comprehension was probed in the context of full sentence structures. Even when sentence length was deliberately kept short, these sentences were rich in semantic content. This inevitably provides a more extensive context than the two word phrases of the current study. Our findings of reduced syntactic comprehension in a contextually deprived context suggest that the availability of additional lexical-semantic information reduces the decline in syntactic comprehension that comes with aging.

The absence of semantic information can be considered an increased processing challenge. In this sense, our interpretation that syntactic decline is more pronounced in the absence of semantic information, is in line with Wingfield, Peelle & Grossman (2003). In this study, the influence of varying processing challenges on syntactic comprehension in older adults was investigated in a different way, by measuring syntactic comprehension of subject-and object relative clause sentences at varying speech rates. While older adults were slower than younger adults at all speech rates tested, this age difference became larger with increased speech rates. In other words, older adults took disproportionally longer to give their comprehension responses at an increased level of processing challenge. Likewise, in the current study, the effect of processing challenge resulted in disproportionately increased response times in older adults when contextual constraints were reduced from a two word phrase with a meaningful content to a similar phrase structure without any representation in the mental lexicon. It should be noted that in the Wingfield, Peelle & Grossman (2003) study, comprehension accuracy only decreased in older adults at very fast speech rates, whereas in the current study, accuracy was already lower compared to young adults for the comprehension of real verb phrases, that is, when processing challenge was at relative minimum. However, as argued above, it could be that the minimal phrases used in the current study already provided a higher processing challenge than the semantically richer sentence structures used by Wingfield, Peelle & Grossman (2003).

This leads us to our third key finding that there was individual variation in the age-related decline in syntactic comprehension. Processing speed provided a unique contribution in explaining the individual variation in performance in the older age group. Increased processing speed was associated with higher performance: older adults with a higher processing speed score were more accurate in comprehending real verb sentences compared to their peers with a lower processing speed score. In addition, in the more challenging pseudoverb block where the older participants as a group showed a significant increase in response time, a higher processing speed score was associated with faster responses. Increased processing speed therefore supported syntactic processing in older adults in two ways: it enabled older adults to be more accurate in their overall faster processing of real verb sentences and to respond faster to the more challenging pseudoverb sentences.

The influence of processing speed on syntactic ability is consistent with a large literature suggesting general processing speed impacts language processing (Waters & Caplan, 2007). Notably, this effect was only present in the older age group in our study. These findings are in line with the contention that the general slowing of processing speed that is associated with age impairs cognitive functioning (Salthouse, 1996). Critically, in the experiment that required the least processing load (the real verb phrases) a faster processing speed decreased the performance gap between older and younger adults.

In addition, the influence of working memory on comprehension performance was different for younger and older adults. For our older adults, a higher working memory capacity was associated with increased comprehension accuracy, irrespective of the lexical semantic context and irrespective of the correctness of the phrase. Furthermore, older adults with a higher working memory capacity experienced a relative advantage in response time in the correct identification of morpho-syntactically correct phrases. These results suggest that, even when the complexity of syntactic processing is reduced to its most basic syntactic operation, increased working memory capacity aids syntactic comprehension in older adults. In the younger age group, the influence of working memory on performance was more limited, emerging only in a subset of the conditions. These findings are in line with Payne et al. (2014) who observed that the effect of working memory on language processing was larger in older compared to younger adults. Our research furthermore demonstrates a similar pattern for processing speed.

However, we are cautious about over-interpreting the observed effects of working memory and processing speed, given that only a single measure was used to assess each cognitive function in this study. The composition of the test battery was aimed at investigating a broad range of common cognitive and physical individual differences. This broad approach is, due to the constraints of potential task fatigue from an expanded additional measurements battery, at the expense of a more in depth measurement of the individual components. To further explore the relationship between comprehension of elementary syntactic structures and these individual components, a more comprehensive assessment by using composite scores consisting of multiple measurements would provide a valuable direction for future research.

In terms of the nature of our syntactic comprehension experiment, it should be noted that both stimuli (two word phrases) and task (grammaticality judgement) were specifically chosen to investigate elementary features of syntactic processing while aiming to maximize the isolation of this process in relation to additional processing mechanisms. As a consequence, certain features related to processing real-life connected speech, such as coarticulatory cues, were either absent or very limited in the decontextualized stimuli of our study. Indeed, compared to processing single words or sentences, processing real-life connected speech has been suggested to rely on additional mechanisms (Alexandrou, Saarinen, Mäkelä, Kujala, & Salmelin, 2017). Moreover, sentence comprehension relies on syntactic processes in a number of ways (Kaan & Swaab, 2002). Therefore, our measure of elementary syntactic comprehension inevitably is a limited proxy of syntactic comprehension more generally. In addition, it should be noted that the differences we observed between young and older adults do not in themselves identify the underlying cause of the effect of age on syntactic comprehension. Age-related effects could, in part, be the result of declines in peripheral and central hearing (Rogers and Peelle, submitted) or auditory-motor speech processing (Panouillères & Möttönen, 2017). However, in our study, accuracy across the board was relatively high for the older adults (specifically, the older adults’ group average accuracy was above 85% in the experimental conditions and even above 90% in the filler conditions). This strongly suggests that participants were able to differentiate correctly among the different experimental conditions, arguing against a profound effect of hearing loss in the present study. Moreover, while older adults were indeed slower and less accurate than younger adults, they were differentially slower and less accurate in response to different experimental manipulations. Therefore, our data pattern cannot be explained in terms of a monotonic effect of slowing or hearing loss due to age.

Another limitation of the current study is that the younger age group consisted of university students, while the older age group was characterised by a more varied educational background. It is possible that this larger variability in the older age group has influenced our findings and may explain our finding that the influence of superior vocabulary on performance was only present in the older age group.

In summary, the results of the current study shed new light on the decline in syntactic comprehension in healthy ageing. Whereas previous studies have primarily investigated complex syntactic structures and focused on syntactic ambiguity, we investigated syntactic comprehension of the elementary building blocks of syntactic processing: syntactic agreement of pronoun and verb. Older adults were slower and less accurate compared to younger adults. This decline seems to increase in the absence of semantic contextual information, which causes older adults to produce slower responses in order to make more accurate decisions. In line with these findings, accuracy for older adults was positively related to processing speed capacity. Taken together, our results provide very clear evidence that syntactic comprehension declines in healthy aging.

## Acknowledgements

We warmly thank all our participants for their contributions to this research and Denise Clissett, the coordinator of Patient and Lifespan Cognition participant database at the University of Birmingham, for recruiting and scheduling participants. We thank Sian Beardsell, Jessica Low, Laura McCue, Zarin Shahabi and Emily Stone for their help with collecting the data. We thank Sophie Hardy and Kelly Garner for their valuable advice on data processing in R and statistical analysis.

## Author contributions

C.P, L.W. and K.S. designed the study. C.P was responsible for data collection and assisted by undergraduate students. C.P. analysed the data. C.P wrote the manuscript under the supervision of L.W. and K.S. All authors revised the manuscript and approved the final version of the manuscript.

## Data Availability

The stimulus materials and the datasets analysed during the current study are available in the OSF repository, [link will be provided upon publication -data can be made available to reviewers].

## Footnotes

1 To preview the results, there was no difference in response bias between the two age groups. A response bias would result in a performance difference between the two conditions. For example, a bias towards responding with ‘yes’ would result in a lower accuracy in the correct syntax condition (‘yes’ here is a mistake) compared to the incorrect syntax condition (‘yes’ here is correct). We ran a t-test to verify whether there was a difference in the mean accuracy between the two conditions for both age groups individually. There was no significant difference in accuracy, neither in the younger age group (t(98) = −0.40, p = 0.69), nor in the older age group (t(98) = 0.12, p = 0.91).

2 However, running the RT model with this outlier participant included did not affect the outcomes.

3 To verify whether the variability in the older age group was larger compared to the younger age group, we performed a Bartlett test between the two age groups for each of the two blocks separately. The results confirm that variability is significantly larger in the older age group, both in the Real Verb block (χ ^2^ (1) = 176.16, p < .001) and the Pseudoverb block χ ^2^ (1) = 20.93, p < .001).

